# Verb-Specific Linking Properties Modulate the N400 Effect: Evidence from Thematic Reversal Anomalies in Malayalam

**DOI:** 10.64898/2026.05.15.725327

**Authors:** S. Shalu, R. Muralikrishnan, Matthias Schlesewsky, Ina Bornkessel-Schlesewsky, Kamal Kumar Choudhary

## Abstract

The present study examined whether thematic reversal anomalies are processed similarly across subject and object experiencer constructions in Malayalam. Event-related brain potentials (ERPs) were recorded as 30 first-language speakers of Malayalam read transitive sentences with the two types of experiencer verbs, in which the thematic role assignment for the preceding arguments was either correct or reverse. The reversal anomaly became apparent only at the position of the experiencer verb. A linear mixed-models analysis confirmed a biphasic N400-P600 effect at the verb for both verb types when the argument roles were reverse. Thus, our results suggest a uniform processing strategy for TRAs irrespective of the type of experiencer verb involved. However, the N400 amplitude was larger for the object experiencer verb compared to subject experiencer verbs. We suggest that the quantitative difference observed for object experiencer verbs is due to the inverse linking of grammatical function and thematic roles associated with these verbs. In other words, verb-specific linking properties modulate the processing of TRAs involving object experiencer verbs. We argue that this modulation occurs because the parser recalibrates cue weighting when the expected form-to-meaning mappings are overridden by the inverse linking properties of object experiencer verbs.

## 1. Introduction

Recent research on language comprehension has increasingly focused on cases where comprehenders form inaccurate or non-veridical representations of the linguistic input (Getty, 2024; Christianson, 2016; Schreiter et al., 2019). Non-veridical language comprehension refers to situations in which the mental representation constructed by comprehenders does not faithfully reflect either the actual linguistic input or real-world plausibility. Rather than always yielding a veridical mapping between sentence structure and meaning, the comprehension system often relies on heuristics or “good-enough” strategies that prioritize efficiency over accuracy. As a result, comprehenders may generate distorted or incomplete interpretations, as seen in garden-path sentences (e.g., initially interpreting “*While Anna dressed the baby played in the crib*” as Anna dressing the baby). Such cases illustrate how non-veridical representations arise naturally in comprehension, offering insights into the cognitive mechanisms that allow humans to interpret language rapidly, even at the cost of accuracy. In this paper, we focus on thematic reversal anomalies (TRAs), a type of non-veridical language phenomenon that has received considerable attention in cross-linguistic sentence processing research, which are known to be influenced by a range of linguistic and methodological factors.

Thematic reversal anomalies arise when the thematic roles assigned to two noun phrases are reversed or swapped within an otherwise syntactically well-formed sentence. In such cases, the surface structure is grammatically correct, yet the sentence is semantically implausible (Kim & Osterhout, 2005; Bornkessel-Schlesewsky et al., 2011). For instance, the sentence “*The fries have #eaten the boys too quickly*” becomes implausible at the verb “*eaten*”, for which *fries* is an acceptable object (as in the case of “*The boys have eaten the fries too quickly*”), but is unsuitable for the subject role, thereby resulting in a linguistic violation (Bourguignon et al., 2012). Such anomalies create a clear mismatch between actual linguistic input and the comprehender’s mental representation of the real event, which ideally should be “*The boys have eaten the fries too quickly*”.

The processing of TRAs has gained significant attention over the years, primarily due to their role in redefining the hitherto assumed rigid alignment in the elicitation of the N400 and the P600 effects with semantics and syntax respectively (Kolk et al., 2003; Kuperberg et al., 2003; Hoeks et al 2004; Kim & Osterhout, 2005; Niewland et al., 2007; Kuperberg, 2007; Bornkessel-Schlesewsky et al., 2011) but also their potential as a valuable tool for investigating linguistic variation (Bornkessel-Schlesewsky et al., 2011). Early studies on TRAs in Dutch and English have consistently observed late parietal positivity effects (the so-called “semantic P600” effects) when compared to plausible control conditions (Kolk et al., 2003; Hoeks et al., 2004; Kim & Osterhout, 2005; van Herten et al., 2005; 2006; Kuperberg et al., 2006, 2007). This finding was novel at the time, as it revealed an unexpected association of P600, traditionally linked to syntactic processing, with semantic processing. However, Schlesewsky and Bornkessel-Schlesewsky’s (2009) study on German TRAs revealed an N400 pattern at the sentence final position, which was in stark contrast to the pattern observed thus far for TRAs. This result called into question the cross-linguistic generalisability of semantic P600s. Several theories have been proposed to account for the P600 and N400 effects observed in response to TRAs (Brouwer et al., 2012), but as discussed in the literature section below, a parsimonious account of all existing findings with respect to TRAs remains elusive to date.

Recent studies have shown that the ERP responses associated with TRAs vary both within and across languages (Bornkessel-Schlesewsky et al., 2011; Bourguignon et al., 2012; Kyriaki et al., 2020) with this variation influenced by factors such as relative prominence of cues to sentence processing (Bornkessel-Schlesewsky et al., 2011; Bornkessel-Schlesewsky & Schlesewsky, 2019), stored semantic relationships (Kolk et al., 2003; Kim and Osterhout et al., 2005; Brouwer et al., 2012; Chow et al., 2016), verb types (Bornkessel-Schlesewsky et al., 2011; Bourguignon et al., 2012; Kyriaki et al., 2020), dialectal differences (Kyriaki et al., 2020), verb dependent selection restrictions (Kuperberg et al., 2007), and various methodological parameters (Chow et al., 2018; Kyriaki et al., 2020; Liao et al., 2022). Among these, verb types have received comparatively less attention, with only a handful of studies explicitly addressing this issue. TRA research on verb medial languages, considering different verb types, has shown that verb-based linking properties (Icelandic, Bornkessel-Schlesewsky et al., 2011) and the thematic aspects related to verb types (English, Bourguinon et al., 2012; Kyriaki et al., 2020) play a crucial role in modulating ERP responses. These findings contrast with accounts that claim the N400 and P600 map directly onto basic lexical processes (Stroud & Phillips, 2012; Brouwer et al., 2012), suggesting instead that they reflect more complex mechanisms involved in sentence comprehension (Kyriaki et al., 2020). These studies have focused on the differences between activity verbs (Act) (e.g; run, cry, etc.) and subject experiencer verbs (SbjExp) (e.g. love, hate, etc.) (Bourguinon et al., 2012; Kyriaki et al., 2020). Although these verbs belong to different semantic classes, they differ only minimally in terms of their configuration involving grammatical function and thematic roles. The Act verbs have a default configuration or direct linking in terms of the hierarchy between grammatical functions and thematic roles (Agent (Subject) > Theme (Object)), whereas the role prototypicality of the arguments is reduced in case of SbjExp verbs (Dowty, 1991; Van Valin & LaPolla, 1997; Primus, 1999; Kyriaki et al., 2020). Nevertheless, the hierarchy of arguments still aligns with the default configuration (Experiencer (Subject) > Stimulus/Theme (Object)). However, verb classes that call for an inversion of the hierarchy have attracted little attention as regards TRA studies. Object experiencer verbs (ObjExp: Experiencer (Object) > Stimulus (Subject)) are a case in point in this regard. To date, this verb class has been employed primarily in investigations related to thematic reanalysis (in comparison to Act verbs) because of their inverse mapping between thematic roles and grammatical functions (Bornkessel et al, 2002; Bornkessel et al., 2003; Bornkessel et al., 2004; Dröge et al., 2014; Gattei et al., 2015). These studies revealed that ObjExp verbs entail higher processing costs compared to Act verbs due to their inverse-linking nature. It remains to be seen, however, how ObjExp verbs are processed in the context of TRAs, given that they have shown distinctive patterns in other contexts.

Although both SbjExp and ObjExp verbs fall within the same semantic class of experiencer predicates, they diverge in how the experiencer roles are grammatically encoded (Croft, 1986; Belletti & Rizzi, 1988; Dowty, 1991; Pesetsky, 1995; Hartshorne et al., 2016; Levin & Grafmiller, 2013) and in their underlying semantic dimensions, including stativity, causality, and volitionality (Hartshorne et al., 2016; Tenny, 1994; Arad, 1998; Landau, 2010). Investigating these verb type contrasts is crucial, as they provide a direct window into how differences in argument structure and semantic properties shape real-time sentence comprehension. Bourguinon et al. (2012) and Kyriaki et al. (2020) tried to include ObjExp verbs in their design and pointed out the challenges in doing so. For instance, TRAs for Act and SbjExp verbs in English are realized at the sentence medial verb, whereas for ObjExp verbs, the TRAs becomes apparent only at the post-verbal object, and therefore the effects of interest are elicited at the object noun instead (refer sentences 1(a) to 1(c)). In other words, at the position when a violation becomes clear for ObjExp verbs, all the arguments of the sentence have been encountered, while for Act and SbjExp verbs only one argument has been encountered when the violation occurs. In a language like English, this makes it difficult to tease apart whether the ERP effect at the ObjExp is due to a violation of the experiencer role, theme role, or both.

1.

a) The answer has #*written* the student on the form.
b) The gifts have #loved the children of the orphanage.
c) The children have pleased the *^#^gifts* of the orphanage.

(Bourguinon et al., 2012)

In view of this, a verb-final language would be ideally suited to examine the processing of TRAs involving ObjExp in comparison to Act and SbjExp verbs. This is because the verb occurs at the end of the sentence in these languages, and therefore regardless of whether the verb is SbjExp or ObjExp or Act, the anomaly would uniformly be realized at the verb position.

To this end, the present study investigated ERP correlates of TRAs involving SbjExp and ObjExp verbs in Malayalam, a verb-final language with rich case morphology. It is important to note that despite extensive research in Indo-European languages, TRAs are yet to be investigated in typologically diverse Indian languages. Therefore, by examining the processing of the TRAs in a Dravidian language such as Malayalam, the present study offers a unique opportunity to test the cross-linguistic validity of existing findings, particularly the effect of verb types.

The remainder of the paper is organized as follows. Subsection 1.1. provides a detailed literature review on the electrophysiology of thematic reversal anomalies. Section 2 outlines the motivation for the present study and our hypothesis. Section 3 describes the methodology, including material used in the study, participant information, and the procedure. Section 4 reports both behavioural and ERP results. Section 5 provides a detailed discussion and interpretation of our findings, and section 6 concludes the paper.

### 1.1. Electrophysiology of Thematic Reversal Anomalies

Previous ERP studies on TRAs across different languages have revealed varying patterns, including monophasic N400 or P600 effects, as well as biphasic N400-P600 patterns. Many theories have been proposed to account for the effects observed in response to TRAs using various single-stream and multi-stream processing models. In the following, we present the literature on TRAs and the interpretation of N400 and P600 ERP components from various perspectives.

Hoeks et al. (2004) found a P600 effect for the sentence “*De speer heft de atleten geworpen* - The javelin has thrown the athletes” in comparison to their control condition “*De speer werd door de atletern geworpen* - The javelin was by the athletes thrown”. They argued that participants experienced some kind of “semantic illusion”, causing them to perceive the sentences as meaningful. However, the semantic illusion persisted only a few hundred milliseconds, after which they quickly detected that there was a problem in the interpretation and engaged in “effortful syntactic processing” in an attempt to revise it. The P600 found in such scenarios has been interpreted as the index of syntactic processing. Hoeks et al. (2004) did not offer an explanation for why the language processor would initiate syntactic revision immediately after interpreting the sentence as though it made perfect sense.

Kim & Osterhout (2005) also found a P600 effect for TRAs such as “The hearty meals were devouring the kids” in comparison to “The hearty meals were devoured by the kids”. They interpreted the P600 in terms of semantic attraction theory, which argued that the evoked response to TRAs is due to the strong semantic attraction between the predicate (e.g. *devoured*) and its argument (e.g. *the hearty meal*). In such cases the semantic information overrides the syntactic information. Thus “*devouring*” in the “*the hearty meal was devouring…*” was said to be interpreted as the wrong inflection of the intended past participle “*devoured*” and this resulted in a P600 effect. However, results from a study by van Herten et al. (2005) provide evidence against the semantic attraction account. The study observed a P600 effect for the sentence “*De vos die op de stropers joeg sloop door het bos* - The fox that hunted[singular] the poachers” in contrast to the sentence “*De stropers die op de vos joegen slopen door het bos* - The poachers that hunted[plural] the fox”. In these sentences both “fox” and “poacher” can possibly “hunt”, but a poacher hunting a fox is more plausible based on world knowledge information, in contrast to a fox hunting a poacher. Further, the number agreement on the verb is correct for the respective subject arguments in this study, unlike the stimuli from Kim & Osterhout (2005). Thus the P600 in van Herten et al. (2005)’s study cannot be attributed to syntactic mismatch between observed and expected verb inflection. They therefore proposed the monitoring theory, with a processing architecture comprising two processing streams: algorithmic stream and plausibility stream. The algorithmic stream is said to handle “syntactic structure building” and the plausibility heuristic stream is responsible for meaningful interpretation based on “lexico-semantic information” and “world knowledge”. According to this account, an N400 effect reflects the relative ease with which the heuristic stream can figure out the interpretation of a sentence. When these two streams produce conflicting interpretations, the processor engages in “monitoring” to identify and resolve the mismatch. This reanalysis process is said to be reflect in a P600 effect.

Kuperberg et al. (2007) observed a P600 for sentences containing TRAs, in which there was no semantic relation between the argument and the verb, as in the sentence “Every morning at breakfast the eggs would watch….” compared to the control condition “Every morning at breakfast the boys would eat toast and jam”. No N400 effect was evoked in their study, but instead a P600 ensued, which contradicts the semantic attraction account, since there is no attraction between “eggs” and “watch”. Further, this result is also difficult to explain in terms of the monitoring theory, since according to the theory, if both the algorithmic stream and the plausibility stream conclude that the sentence does not make sense, the parser should evoke an N400, and no P600 effect, since there is no conflict between the two processing streams. To address this, Kuperberg et al. (2007) proposed “continued combinatory analysis”, involving three parallel streams: (1) the semantic memory-based stream draws plausibility information based on the “lexico-semantic information” and “world knowledge information” from the long-term memory; (2) the syntax driven stream controls the way words combine together based on morpho-syntactic information; and (3) the thematic role based stream assigns thematic roles to the arguments based on lexical, thematic cues such as animacy. They argued that the latter two streams work in a combinatory manner. Whenever the syntax-driven stream comes into conflict with the thematic-role based stream, the parser engages in what is termed a “continued combinatory analysis” – a revision process aimed at resolving the conflict between the two streams. This revision process is typically reflected in the elicitation of a P600 effect. This is for instance the case when the syntactic stream assigns an inanimate NP (*the eggs*) the role of actor, while the thematic stream assigns it the role of undergoer instead. In such cases, an N400 effect is typically absent, as the conflict between the combinatory streams is thought to hinder further semantic processing within the memory-based stream. However it is unclear how this account could explain the biphasic N400-P600 pattern reported in Kuperberg et al. (2010) for sentences such as “The journalist astonished the article before his coffee break” in comparison to “The journalist wrote the article before his coffee break”.

Schlesewsky and Bornkessel-Schlesewsky (2009) employed TRAs in German and observed a biphasic N400-P600 pattern at the sentence final position for irresolvable anomalies, but only an N400 effect for resolvable anomalies. This was different from the pattern of results reported thus far for thematic reversal anomalies. Similarly Bornkessel et al. (2011) also found biphasic N400-P600 effects for Icelandic but a monophasic N400 for TRAs in both Turkish and Chinese. They interpreted these results in terms of their extended Argument Dependency model (eADM; Bornkessel-Schlesewsky & Schlesewsky, 2006; 2008, 2009; 2014; 2016). As per this model, Bornkessel et al. (2011) argued that in languages that rely on word order to identify argument roles such as Dutch, English and Icelandic (in the case of alternating verbs), a processing problem is realized only in the generalized mapping step when the compute linking and plausibility processing steps produce noncompatible results and evoke P600 effects. Conversely, in languages that primarily use other prominence cues like case marking, animacy, or other grammatical features to determine argument roles, such as Chinese, German, Icelandic (non-alternating verbs) and Turkish, the problem already occurs in the compute linking stage (which links role assignments to the verb’s logical structure, i.e., its decomposed semantic representation), which engenders an N400 effect. The P600 effect observed in both sequence-dependent and sequence-independent languages has been interpreted as a late, target-related P300, reflecting a binary categorization of the stimulus as well-formed or ill-formed. It is typically observed when the grammar of the language permits a clear binary decision — anomaly versus non-anomaly — at the position of the critical word.

The eADM accounts for much of the variability observed in the processing of TRAs discussed above. However, the N400 effect reported by Kos et al. (2010) for the sentence *“Fred eats a restaurant”* would be contrary to the predictions of this model. According to the eADM, the compute prominence would assign *“a restaurant”* the role of undergoer, while the plausibility processor may reinterpret it as a locative (e.g., *eating in a restaurant*). Since both processing streams yield a coherent interpretation, the model predicts no N400 effect. Instead, any conflict should emerge only at the generalized mapping stage, potentially eliciting a P600. Contrary to this prediction, Kos et al. observed a robust N400 and no P600. To explain this Kos et al. (2010) proposed the Processing Competition Model, which posits two parallel processing streams - syntactic and semantic - each independently constructing interpretations of the incoming material. When the outputs of these streams conflict, the resolution burden falls on the stream with a weaker contextual support. If semantic cues dominate, the syntactic stream resolves the conflict, resulting in a P600. Conversely, if syntactic cues are stronger, the semantic stream bears the burden, leading to an N400. In the sentence “Fred eats a restaurant”, the syntactic stream produces an implausible analysis (Fred consuming a restaurant), while the semantic stream offers a plausible one (Fred eating in a restaurant). Kos et al. (2010) argue that syntactic cues are stronger here because they clearly predict a noun phrase (NP), unlike semantic cues which lack a specific expectation. As a result, the semantic system bears the burden of resolving the conflict, leading to an N400 effect, but no P600. In contrast, in a sentence like “The javelin has thrown the athletes”, syntax yields an implausible reading, while semantics supports a plausible one (athletes throwing the javelin). Here, semantic cues are stronger, shifting the conflict resolution to the syntactic stream, resulting in a P600 effect, but no N400. Despite this explanatory power, the Kos et al. (2010) model still struggles to account for biphasic N400–P600 effects.

After a comprehensive review of various multi-stream models for the P600 and N400 effects found for the TRAs, Brouwer et al. (2012) concluded that these models are unable to explain the full range of evidence for N400 and P600 effects observed in cross-linguistic investigation. Brouwer et al. (2012) proposed the “Retrieval-Integration” account. The model advocates for a single stream architecture for sentence comprehension and does not see a need for a separate stream for semantic processing. According to this model, the N400 and the P600 index two consecutive stages of processing, namely retrieval and integration. The N400 amplitude is linked to the retrieval phase, during which all the information tied to an incoming word – such as its syntactic, semantic, and pragmatic features - is accessed from long-term memory (Kutas and Federmeier, 2000, 2011; Lau et al., 2008; 2009; Federmeier and Laszlo, 2009; van Berkum, 2009, 2010). Importantly, the N400 does not reflect the process of combining or integrating this information into the sentence context. The integration of this activated lexical information into the current mental representation of an unfolding sentence is indexed by the P600 amplitude. For instance, the sentence “the javelin has thrown the athletes” evoked a P600 effect compared to the control condition “The athletes has thrown the javelin”. There is no retrieval issue in terms of the retrieval-integration account here, since lexical information associated with “thrown” is the same in both the sentences due to word and context priming. For the reversal sentence there will be difficulty in integrating this information into the existing mental representation, hence a P600 effect is evoked. Integration is predicted to be difficult because it results in a representation that contradicts what we know about the world: javelins are not able to, and do not usually throw athletes. In the sentence “The javelin has the athletes summarized”, there is no semantic or lexical priming for the verb “summarized” in the context of the noun “javelin”. Since a javelin cannot possibly summarize anything, the brain has a harder time retrieving the lexical/semantic features of that verb. This difficulty leads to a larger N400 effect. The recent discussion by Delogu et al. (2025) provides strong support for a single-stream language processing model, such as the retrieval-integration framework proposed by Brouwer et al. (2012). They interpret the biphasic ERP pattern often observed in response to linguistic anomalies as reflecting increased processing effort during both the retrieval and integration stages. When both of these processes demand substantial cognitive effort, a P600 component is typically elicited. However, when retrieval and integration processes overlap significantly in time, the P600 may be weakened or even be absent from the ERP data. Notably, deeper levels of comprehension are thought to enhance the P600 response, making it more detectable even when it overlaps with the N400. A recent finding from Chinese TRAs also supports the retrieval-integration based interpretation for the N400 and P600 effects (Zhang et al., 2025).

However findings from Bourguignon et al. (2012) cannot be accounted for by the retrieval-integration account, as they found an N400-P600 effect for TRAs involving SbjExp verbs, and a P600 effect for TRAs involving action verbs. As there is no retrieval difficulty in either case, and lexical information associated with TRAs involving verbs like “eaten” and “love” in both the TRA conditions and their control condition counterparts are equally accessible in both sentence types, and therefore no N400 should ensue according to the retrieval-integration account. However, an N400 indeed ensued for reversal anomalies involving SbjExp verbs (Bourguignon et al. 2012). Besides the verb type based differences in ERPs, the N400 effect is novel for English in the context of TRAs, as previous studies have only reported a P600. The presence of an N400 effect in response to role reversal in English also contradicts the Bornkessel-Schlesewsky et al. (2011) account, which posits that the N400 effect is influenced by whether verb-argument linking is sequence-dependent or sequence-independent.

Bourguignon et al. (2012) interpreted the verb dependent N400 effect in terms of the aspectual-thematic structure between the two verb types based on the eADM model. In the case of SbjExp verbs, the prototypical role assigned by the compute prominence step (assigning actor-undergoer) needs to be reanalysed as experiencer at the compute linking step (analysing thematic relationship based on the verb’s logical structure). The N400 effect found in the study is then said to be an index of this reanalysis. The P600 effect found in the study is smaller than in previous studies investigating TRA, which may be due to the influence of sentential context, working memory capacity of participants and the salience of the items. However, such an interpretation does not account for the data reported in Kyriaki et al. (2020) and Ehrenhofer et al. (2022). Kyriaki et al. (2020) replicated the Bourguignon et al. (2012) study by varying the modality and tasks, and found N400 effects for both verb types (albeit in differing task and modality combinations).

Kyriaki et al. (2020) speculated that the verb-independent N400 effect observed in their study could be due to the dialectal differences in the varietyy of English use in their study, because previous research on TRA in English has largely focused on North American English, whereas Kyriaki et al. (2020) examined Australian English. However, the absence of a P600 effect for TRAs involving SbjExp verbs was unexpected compared to Bourguignon et al., (2012). They argued that it is because of the decreased salience of the violation involving subject experiencer verbs compared to violations in their filler sentences, which led to a diminished ERP response. Overall, the study suggests that ERP effects in response to TRA constructions are sensitive to various methodological factors.

Table 1 provides an overview of cross-linguistic ERP studies that examined thematic reversal anomalies and the ERP components they reported. Despite this body of work, a unified explanation that parsimoniously accounts for all cross-linguistic ERP findings on TRAs remains elusive. The present study seeks to examine the phenomenon employing experiencer verbs in Malayalam, and thereby complements existing findings from a typologically different language compared to languages hitherto investigated. As mentioned earlier, the effect of verb types on reversal anomaly is less explored, and to the best of our knowledge, ObjExp verbs have not been examined in this regard. Hence, we employ SbjExp and ObjExp verbs in the present study and test the cross-linguistic generality of these effects, particularly with respect to studies which have explored the effect of verbs types in the context of TRAs.

**Table 1:**
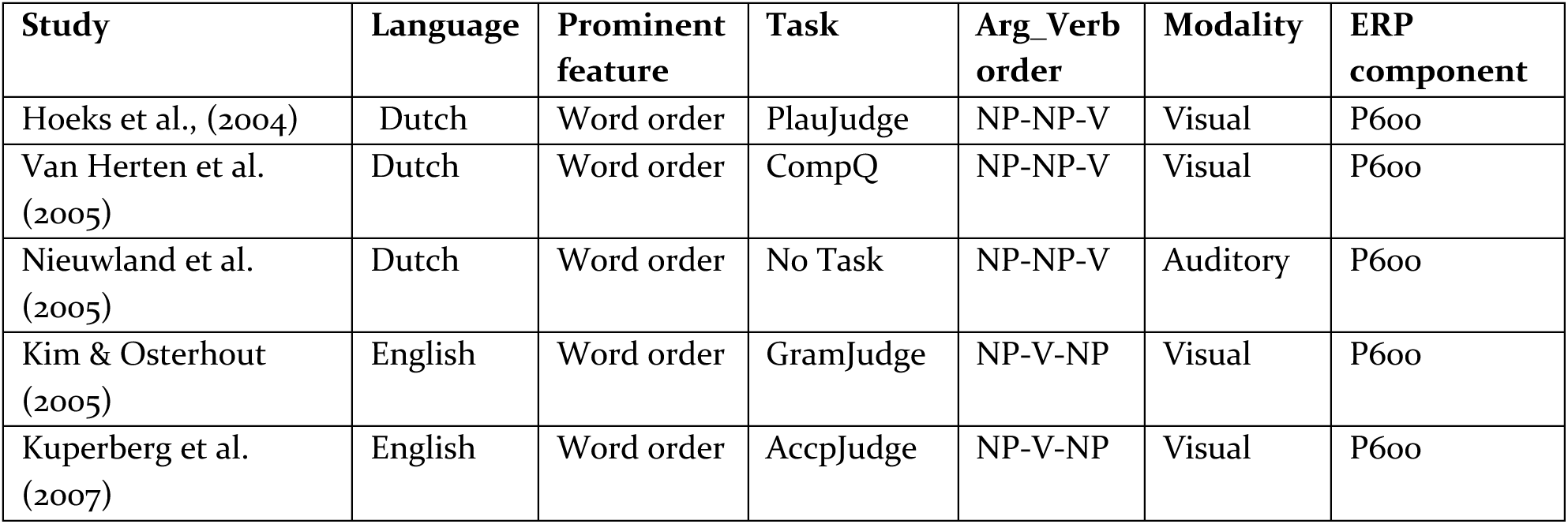
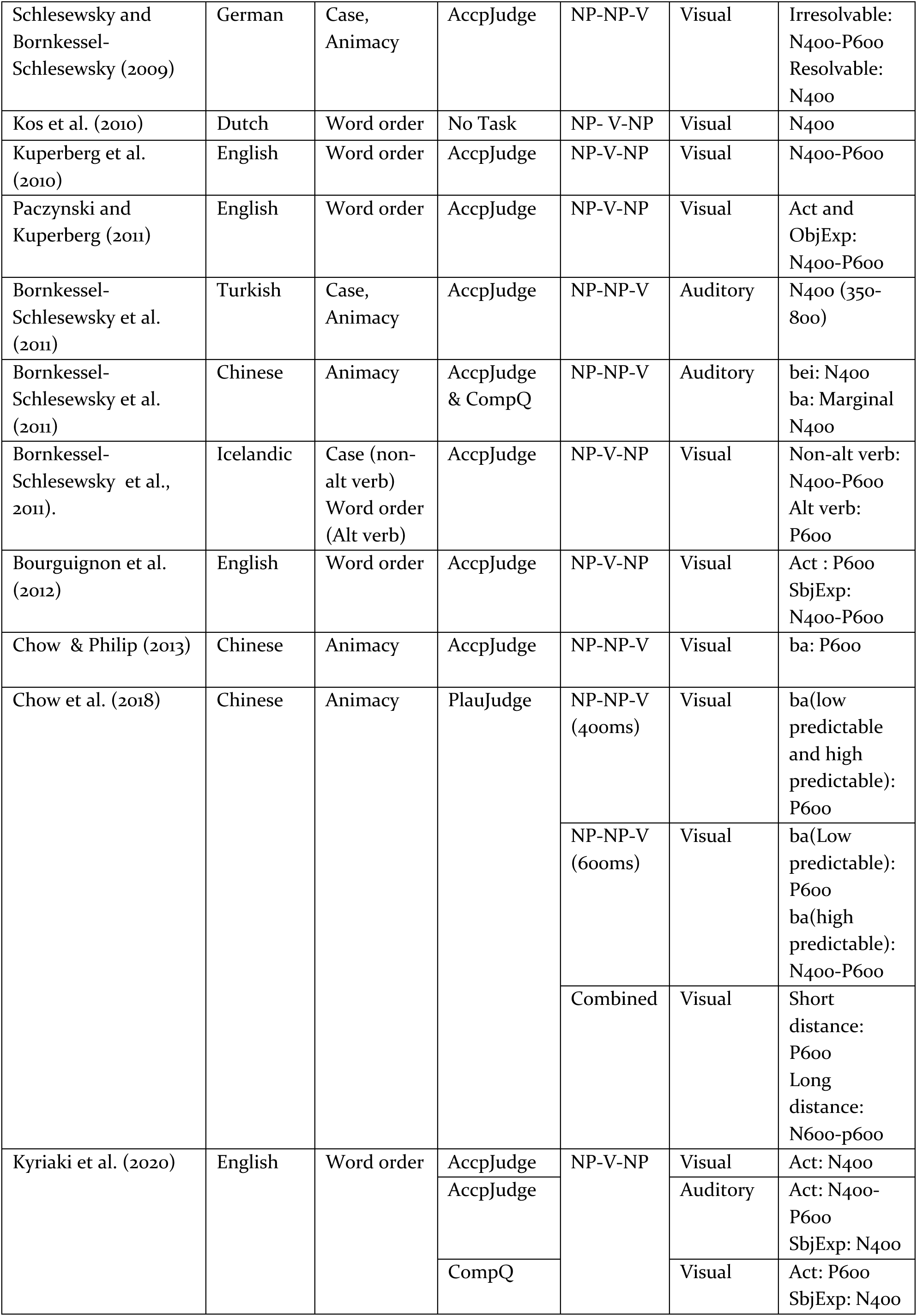
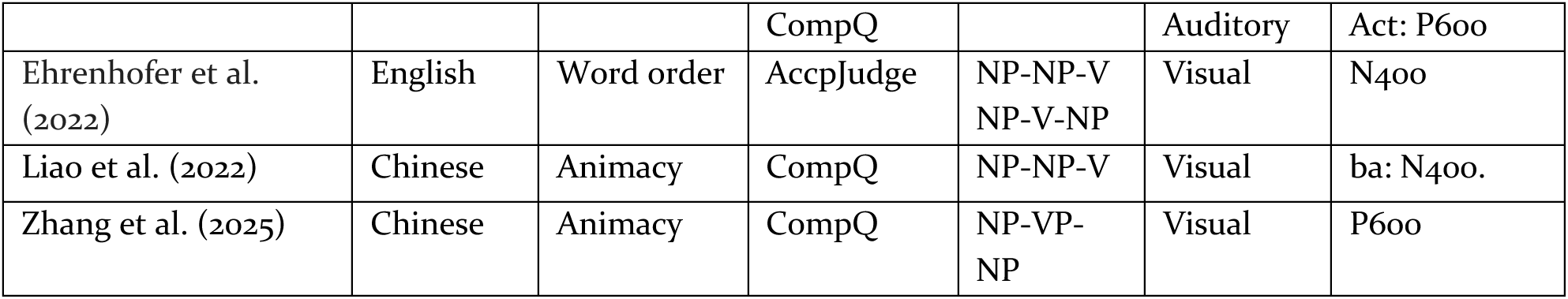
An overview of cross-linguistic ERP studies that examined thematic reversal anomalies. The comprehension question task is abbreviated as CompQ and the acceptability judgement task as AccpJudge.

| Study | Language | Prominent feature | Task | Arg_Verb order | Modality | ERP component |
| --- | --- | --- | --- | --- | --- | --- |
| Hoeks et al., (2004) | Dutch | Word order | PlauJudge | NP-NP-V | Visual | P600 |
| Van Herten et al. (2005) | Dutch | Word order | CompQ | NP-NP-V | Visual | P600 |
| Nieuwland et al. (2005) | Dutch | Word order | No Task | NP-NP-V | Auditory | P600 |
| Kim & Osterhout (2005) | English | Word order | GramJudge | NP-V-NP | Visual | P600 |
| Kuperberg et al. (2007) | English | Word order | AccpJudge | NP-V-NP | Visual | P600 |
| Schlesewsky and Bornkessel-Schlesewsky (2009) | German | Case, Animacy | AccpJudge | NP-NP-V | Visual | Irresolvable: N400-P600<br>Resolvable: N400 |
| Kos et al. (2010) | Dutch | Word order | No Task | NP- V-NP | Visual | N400 |
| Kuperberg et al. (2010) | English | Word order | AccpJudge | NP-V-NP | Visual | N400-P600 |
| Paczynski and Kuperberg (2011) | English | Word order | AccpJudge | NP-V-NP | Visual | Act and ObjExp: N400-P600 |
| Bornkessel-Schlesewsky et al. (2011) | Turkish | Case, Animacy | AccpJudge | NP-NP-V | Auditory | N400 (350-800) |
| Bornkessel-Schlesewsky et al. (2011) | Chinese | Animacy | AccpJudge & CompQ | NP-NP-V | Auditory | bei: N400<br>ba: Marginal N400 |
| Bornkessel-Schlesewsky et al., (2011). | Icelandic | Case (non-alt verb)<br>Word order (Alt verb) | AccpJudge | NP-V-NP | Visual | Non-alt verb: N400-P600<br>Alt verb: P600 |
| Bourguignon et al. (2012) | English | Word order | AccpJudge | NP-V-NP | Visual | Act : P600<br>SbjExp: N400-P600 |
| Chow & Philip (2013) | Chinese | Animacy | AccpJudge | NP-NP-V | Visual | ba: P600 |
| Chow et al. (2018) | Chinese | Animacy | PlauJudge | NP-NP-V (400ms) | Visual | ba(low predictable and high predictable): P600 |
|  |  |  |  | NP-NP-V (600ms) | Visual | ba(Low predictable): P600<br>ba(high predictable): N400-P600 |
|  |  |  |  | Combined | Visual | Short distance: P600<br>Long distance: N600-p600 |
| Kyriaki et al. (2020) | English | Word order | AccpJudge | NP-V-NP | Visual | Act: N400 |
|  |  |  | AccpJudge |  | Auditory | Act: N400-P600<br>SbjExp: N400 |
|  |  |  | CompQ |  | Visual | Act: P600<br>SbjExp: N400 |

|  |  |  | CompQ |  | Auditory | Act: P600 |
| --- | --- | --- | --- | --- | --- | --- |
| Ehrenhofer et al. (2022) | English | Word order | AccpJudge | NP-NP-V<br>NP-V-NP | Visual | N400 |
| Liao et al. (2022) | Chinese | Animacy | CompQ | NP-NP-V | Visual | ba: N400. |
| Zhang et al. (2025) | Chinese | Animacy | CompQ | NP-VP-NP | Visual | P600 |

## 2. The Present Study

The main goal of this study, as outlined earlier, is to examine the difference between SbjExp and ObjExp verbs in terms of their neural response to TRAs in Malayalam. We devised a 2 x 2 factorial design involving transitive constructions and manipulated the thematic role assignment (correct versus reverse) by employing different animacy combinations of subject and object arguments entailing correct versus reversed thematic assignments, which in turn depended upon the verb type^1^ (SbjExp versus ObjExp verbs). First-language speakers of Malayalam read transitive sentences of the form NP1-Nom—NP2-Acc—Verb (Table 2), in which the thematic role assignments of NP1 and NP2 (correct versus reverse) became clear only at the position of the verb based on whether the verb was a SbjExp or an ObjExp verb. That is, all subject and object combinations in our stimuli were felicitous and well-formed until the verb was encountered.

**Table 2.** An example set of experimental stimuli. In the condition labels, the experiencer verb type is indicated as SbJExp (subject experiencer verb) and ObjExp (object experiencer verb); the thematic roles are indicated Correct (role assignments are correct) and Reverse (role assignments are reversed). The conditions SbjExp_Correct and ObjExp_Correct constitute acceptable sentences with correct thematic role assignments, and the conditions SbjExp_Reverse and ObjExp_Reverse constitutes thematic reversal anomalies.

| Condition | Sample Stimulus |
| --- | --- |
| SbjExp_Correct | Subeena sammanathe snehichu.<br>Subeena (An.F.Nom) gift (In)-Acc love-Pst<br>“Subeena loved the gift.” |
| SbjExp_Reverse | #Sammanam Subeenaye snehichu. |

**Table 2.** An example set of experimental stimuli.
|  |  |
| --- | --- |
|  | Gift (In.Nom) Subeena (An.F)-Acc love-Pst<br>“Gift loved Subeena.” |
| ObjExp_Correct | Sammanam Subeenaye ahaldahippichu.<br>Gift (In.Nom) Subeena (An.F)-Acc delight-Pst<br>“The gift delighted Subeena.” |
| ObjExp_Reverse | #Subeena Sammanathe ahladhippichu.<br>Subeena (An.F.Nom) gift(In)-Acc delight-Pst<br>“Subeena delighted the gift.” |

Since there is no published TRA study on Dravidian languages to date, we have drawn on processing literature from other languages that have examined TRAs – particularly those that are verb-final and case-marked – as well as theoretical literature on experiencer verbs to formulate our hypothesis. This is motivated by the fact that Malayalam is morphologically rich with respect to case-marking, and exhibits a canonical verb-final structure. Based on reversal studies on German, Turkish (Bornkessel-Schlesewsky et al., 2011) and reversal studies that employed experiencer verbs (Bourguignon et al., 2012; Kyriaki et al., 2020), we hypothesize that the TRAs involving both the verb types (SbjExp and ObjExp verbs) should evoke qualitatively similar neural correlates compared to their control counterparts (non-TRA), since both predicates have a similarly special status, i.e., both are psychological-experiencer predicates (Bornkessel et al., 2003). A further similarity between the reversal conditions is that, though the reversal condition results in the experiencer role being assigned to the inanimate subject in case of SbjExp and to the inanimate object in case of ObjExp, in both cases, the verbs however do not permit the experiencer argument to be a non-sentient entity, leading to a strong violation of animacy constraints. Therefore, we predict an N400 effect for TRAs involving both verb types, in line with previous studies on verb-final languages such as Turkish and German (Schlesewsky and Bornkessel-Schlesewsky et al., 2009; Bornkessel-Schlesewsky et al., 2011). Additionally, we expect a P600 effect for both reversal sentences, as they are irresolvable and do not constitute a structural ambiguity at the critical position (verb) at which the reversal anomaly becomes apparent (Bornkessel-Schlesewsky et al., 2011). That is, we expect to observe an N400–P600 pattern for both types of reversal sentences. Our predictions are in line with findings from Bourguignon et al. (2012), who reported an N400–P600 pattern in response to reversal anomalies involving SbjExp verb, and partially with findings from Kyriaki et al. (2020).

Alternatively, since ObjExp verbs are more complex than SbjExp verbs in terms of both syntax and semantics, TRAs involving SbjExp and ObjExp verbs should show different processing mechanisms compared to their respective control counterparts. This would be in line with previous studies that found that verb types modulate the ERP components elicited by thematic reversal anomalies (Bornkessel-Schlesewsky et al., 2011; Bourguignon et al., 2012; Kyriaki et al., 2020). Thematic reversals involving SbjExp and ObjExp verbs differ in several important ways. As outlined in the Introduction, these two verb classes differ significantly in how grammatical relations are linked to thematic roles. SbjExp verbs follow a direct linking pattern, whereas ObjExp verbs exhibit an inverse linking pattern (Wilson & Dillon, 2022). Hence, TRAs involving ObjExp verbs do not simply reverse roles in the same way as those involving SbjExp verbs. Rather, they introduce a second layer of disruption on top of an already non-canonical linking inherent to ObjExp verbs. That is, in ObjExp_Reverse, two hierarchy inversions occur simultaneously: (i) the inverse linking of grammatical functions to semantic roles intrinsic to ObjExp verbs and (ii) the reversal-induced violation of verb-specific role expectations (which is, in effect, the reversal of the intrinsic hierarchy inversion). That is, in contrast to SbjExp reversals that involve a single disruption of otherwise canonically aligned hierarchies, ObjExp reversals involve a dual mismatch, requiring the comprehender to resolve both the role reversal and the intrinsic hierarchy inversion. Consequently, the TRAs involving ObjExp verbs are predicted to be more difficult to process than TRAs involving SbjExp verbs. In addition to these differences, the two verb classes diverge along several other semantic dimensions as well. SbjExp are typically associated with a greater degree of volition, non-causality, and often encode mental experiences, whereas ObjExp verbs are more often characterized by reduced volitionality, stronger causal relations, and mostly encode physical experiences (Brennan and Pylkkanen, 2010; Brennan, 2015; Dowty, 1991; Croft, 1986; Tenny, 1994; Arad, 1998; Landau, 2010; Hartshorne et al., 2016). Based on the accounts of Bornkessel-Schlesewsky et al. (2011), Bourguignon et al. (2012) and Kyriaki et al. (2020), one straightforward prediction is that there may be an amplitude difference in the N400 component for TRAs between the two verb types. Another potentially relevant prediction derives from the finding of a LAN effect at the position of a clause-final ObjExp in subject-before-object sentences in German (Bornkessel et al., 2004), interpreted as reflecting the mismatch between grammatical function (Subject > Object) and thematic role hierarchies (Experiencer > Theme). However, it remains to be seen whether and in what manner this predicted difference between the subject experiencer and object experiencer verbs ensues.

## 3. Method

### 3.1. Ethical Statement

The research protocol for this experiment was approved by the Institute Ethical Committee (IEC) of the Indian Institute of Technology Ropar, where the experiment was conducted. Informed and written consent was obtained from participants when they arrived at the lab. All the assessments thus carried out were in accordance with the guidelines and regulations approved by the committee.

### 3.2. Participants

3o first-language speakers of Malayalam (mean age = 27.7; age range = 18 - 40; 16 females; 14 males), mostly students and staff at the Indian Institute of Technology Ropar, living in Ropar, India, participated in the experiment. Each participant was remunerated for their participation as per the allowance permitted by the ethical committee. Before the start of the experiment, the participants were briefed about the experiment and informed consent was obtained from them for the use of their data for academic purposes. All the participants were right-handed as evaluated by an adapted version of the Edinburgh Handedness Inventory (Oldfield, 1971) in Malayalam. The participants had normal or corrected-to-normal vision and had no known neurological disorder at the time of their participation in the experiment. All the participants were first-language speakers of Malayalam and reported having acquired the language before the age of six. As is the case with most of the Indian population, all of our participants spoke one or more additional languages. Data from 6 further participants were not included for analysis due to excessive artefacts.

### 3.3. Materials

We employed 10 SbjExp verbs and 10 ObjExp verbs for constructing the critical conditions in our study and each verb was repeated 3 times. Additionally, we used 15 different inanimate object nouns, each repeated twice across critical conditions and used 30 different names as our subject nouns. This design resulted in 30 sets of sentences across four critical conditions, as shown in Table 1, yielding a total of 120 critical sentences (see supplementary material). Subject nouns were always in nominative case and object nouns were always accusative case marked. Of the 120 critical sentences, 60 sentences were implausible due to role reversal of the arguments, and 60 sentences were plausible. Additionally, we constructed 480 filler items to introduce variety in the stimuli to avoid strategic responses from the participants. These fillers were interspersed with the critical items such that there was a total of 600 sentences. These items were pseudorandomized for presentation during the experiment.

### 3.4. Procedure

The experiment consisted of a practice session followed by the actual experiment, with all the activities, including electrode preparation and stimulus presentation lasting approximately 3 hours. Initially, the procedure and the tasks to be performed during the experiment were explained to the participants, and a printed instruction sheet was provided to them. However, the question under investigation in the experiment was not revealed to the participants for the acquisition of unbiased data. Then, they filled a consent form giving informed consent for their participation along with a Malayalam version of the Edinburg handedness questionnaire to evaluate their handedness. Finally, the head measurements of the participants were taken, and a Hydrocel GSN net of the appropriate size was placed on their scalp. They were seated within a soundproof chamber on a comfortable chair at a distance of 1 m from a 20” LCD monitor on which the stimuli were presented using E-prime 2.0 (Psychology Software Tools, Pittsburgh, PA) (https://pstnet.com).

The structure of each experimental trial was as follows. Each trial began with a fixation ‘+’ sign at the centre of the screen for a period of 1000 ms. This was followed by a blank screen for 100 ms. Then, the stimulus sentence was presented using the Rapid Serial Visual Presentation (RSVP) method, with each sentence (including both critical and filler trials) shown in three chunks. Each chunk appeared on the screen for 700 ms, followed by a 100 ms inter-stimulus interval (ISI). This pace ensured comfortable reading for participants and aligns with existing ERP research on orthographically and morphologically complex languages such as Arabic (Muralikrishnan & Idrissi, 2021), Mandarin Chinese (Wang et al., 2009), Icelandic (Bornkessel-Schlesewsky et al., 2020), Japanese (Wang & Schumacher, 2013), Turkish (Demiral et al., 2008), as well as prior ERP studies on Malayalam (Shalu et al., 2025, 2026). Following the last chunk, participants performed two tasks. Given the use of a violation paradigm, the first was an acceptability judgment task: when prompted by a “???” sign on the screen, participants pressed a green button if they considered the sentence acceptable, and a red button if not. To ensure attentive reading, this was followed by a probe task. After completing the acceptability judgment task or when 1500 ms had passed, a probe word appeared on the screen for 2000 ms. Participants had to press the green button if the word had appeared in the preceding sentence, or the red button if it had not. Half of the probe words were present in the previous trial and required a “yes” response; the remaining half were novel and required a “no” response. The placement of green and red response buttons was counterbalanced across participants. For half of them, the ‘yes’ response was mapped to the left key and the ‘no’ response to the right key, while the other half had the opposite configuration. Participants were instructed to refrain from blinking during the presentation of words but were allowed to blink while completing the two response tasks. Prior to the main experiment, participants completed a practice session consisting of 18 items to become familiar with the trial structure. These practice items did not include any stimuli from the main experiment. The experimental session itself was divided into 15 blocks, each containing 40 sentences, with brief rest periods between blocks. Upon completing the experiment, participants were asked to fill out a questionnaire regarding their experience.

### 3.5. EEG Recording, Pre-processing and Statistical Analysis

The scalp EEG was recorded using 32 Ag/AgCI electrodes fixed on the scalp using the Hydrocel Geodesic Sensor Net, with Cz serving as the online reference. The electrooculogram(EOG) was recorded by means of electrodes placed at the outer canthi and under the eyes for horizontal and vertical eye movements respectively, and the interelectrode impedance was kept below 50 kΩ (amplifier input impedance > 1 GOhm) as per system recommendations (Ferree et al., 2001). All EEG and EOG signals were amplified using a Net Amps 400 Amplifier. The data was recorded at a sampling rate of 500 Hz.

The EEG data was pre-processed for further analysis using the EEGLAB toolbox (Version 14; Delorme & Makeig, 2004, sccn.ucsd.edu) in MATLAB (Version R2023b; The MathWorks, Inc.). The data was down-sampled to 250 Hz and filtered using a 0.3–20 Hz band-pass filter in order to remove slow signal drifts. These filter settings are sufficiently broad to include language-related ERP activity that is typically in the frequency range of about 0.5–5 Hz (Delorme, 2023; Roehm et al., 2002), and have been employed in several previous cross-linguistic ERP studies on language processing in diverse languages. This data was then re-referenced offline to the average of the two mastoids, and channels on the outer extent of the face and head were removed. In a 1 Hz high-pass filtered copy of this original data, an Independent Component Analysis (ICA, Iriarte et al., 2003) was computed. Bad channels were removed in this data before submitting it to an ICA computation using the extended Infomax algorithm. The resulting Independent Components (ICs) were then tested using the ICLabel plugin (Pion-Tonachini et al., 2019) for identifying and marking ICs that were artefactual. The weights computed during ICA were then copied to the original data, and the ICs marked as artefactual were rejected from the original data. The bad channels removed before computing the ICA were interpolated from the remaining channels. The data was then imported in R (Version 4.4.2; R Core Team, 2024) using the eeguana package (Version 0.1.11.9001; Nicenboim, 2018) for epoching and statistical analysis.

Data epochs were extracted from the continuous data for each participant for the critical conditions at the position of the verb, from 200 ms before the onset until 1200 ms after the onset of the verb (i.e., −200 to 1200 ms). Epochs in which the amplitude exceeded the threshold of 100 μV in either direction were rejected, as well as those in which the difference between the minimum and maximum amplitudes within a window of 200 ms crossed the threshold of 100 μV. Furthermore, trials in which the acceptability judgement task was not performed were also rejected. Data from participants with too few remaining trials were excluded from further analysis. For a given participant, data epochs from about 25 trials remained in each condition after these rejections, and therefore the number of valid trials entering the analysis did not differ much across conditions. Across participants, there were a total of 3076 data epochs / trials entering the analysis, with about 769 trials per critical condition. For the purposes of visualization, the valid epochs were averaged across items per condition per participant, and then the grand-averages were computed across participants and smoothed using an 8 Hz low-pass filter to obtain ERP plots at the verb for each condition.

#### 3.5.1. ERP Data Analysis

The single trial EEG epochs at the verb for each critical condition were used for statistically analysing the mean amplitudes in selected time-windows of interest by fitting linear mixed effects models using the lme4 package (Version 2.0.0, Bates et al. 2015) in R (Version 4.5.2, R Core Team, 2025). Interactions were resolved by computing estimated marginal means on the response scale using the emmeans package (Lenth & Piaskowski, 2025) with the mvt multivariate adjustment applied for multiple comparisons. The statistical models included the fixed factors Verb type (SbjExp and ObjExp) and Thematic roles (Correct and Reversed), as well as the topographical factor Regions of Interest (ROI). The ROIs were defined by clustering topographically adjacent electrodes in 6 lateral and 2 midline regions. The lateral ROIs were: Left-Frontal, comprised of the electrodes E3 and E11 (equivalent to F3 and F7 in the 10-20 electrode system); Left-Central, comprised of the electrodes E5 and E13 (C3 and T7); Left-Parietal, comprised of the electrodes E7 and E15 (P3 and P7); Right-Frontal, comprised of the electrodes E4 and E12 (F4 and F8); Right-Central, comprised of the electrodes E6 and E14 (C4 and T8); and Right-Parietal, comprised of the electrodes E8 and E16 (P4 and P8). The midline ROIs were: Mid-Fronto-Central, comprised of E17 and E28 (Fz and ∼FCz); and Mid-Parieto-Occipital, comprised of E19, E20, E9 and E10 (Pz, Oz, O1 and O2). Furthermore, rather than performing a traditional subtraction-based baseline correction, the mean 200 ms pre-stimulus baseline amplitude (−200 to 0 ms) in each data epoch was included as a covariate (after scaling) in the statistical model. This was done in order to account for and regress out potential contributions of differences in the baseline period in the statistical analysis (Alday, 2019). However, we do not interpret effects involving pre-stimulus amplitudes, because these did not form part of our hypotheses. Further, this is in line with the fact that “we can include additional covariates as controls without further interpreting those covariates” (Alday, 2019, p.9). The contrasts for the categorical factors used sum contrasts (scaled sum contrasts for two-level factors), such that coefficients reflect differences to the grand mean (Schad et al., 2020). Except where specified otherwise, the random effects structure was maximal (Barr et al., 2013), with random intercepts for participants and items, and by-participant random slopes for the effect of Verb type, Thematic roles and their interaction term. A by-item random slope specification was not included because it would not be identifiable for this data: the items were coded uniquely across conditions such that they indicated a combination of condition + item. Following modern statistical recommendations, we do not use the term ‘statistically significant’ or its variants based on p-value thresholds (Wasserstein et al., 2019), but report precise p-values as continuous quantities (e.g., p = 0.06 rather than p < 0.08), unless a value is “below the limit of numerical accuracy of the data”, in which case, we report it as p < 0.001 (Amrhein et al., 2019, p.206). Further, we supplement the p-values by transforming them into s-values (Shannon information, surprisal, or binary logworth) and report s = – log_2_(p), which provides a nonprobability measure of the information provided by a p-value on an absolute scale (Shannon, 1948; Greenland, 2019). In other words, “the s-value provides a gauge of the information supplied by a statistical test” and has the advantage of providing “a direct quantification of information without” requiring prior distributions as input (Rafi & Greenland, 2020, p.6).

## 4. Results

### 4.1. Behavioural Data

The mean acceptability ratings as well as the probe detection accuracy for the critical conditions, shown in Table 3, were calculated for the trials that entered the ERP analysis using the behavioural data collected during the experiment. As mentioned earlier, only those trials for which the acceptability judgement task was performed (i.e., not timed out) were considered for analysis. Acceptability was highest for the correct conditions, whereas it was lowest for the reverse conditions. The probe detection accuracy was very high across all conditions. Figure 1 shows raincloud plots (Allen et al., 2021) of the behavioural acceptability judgements. Panel A shows the by-participant variability of acceptability ratings, with the individual data points representing the mean by-participant acceptability of each Verb type and Thematic roles combination. Panel B shows the by-item variability of acceptability ratings, with the individual data points representing the mean by-item acceptability of each Verb type and Thematic roles combination.

**Figure 1.**
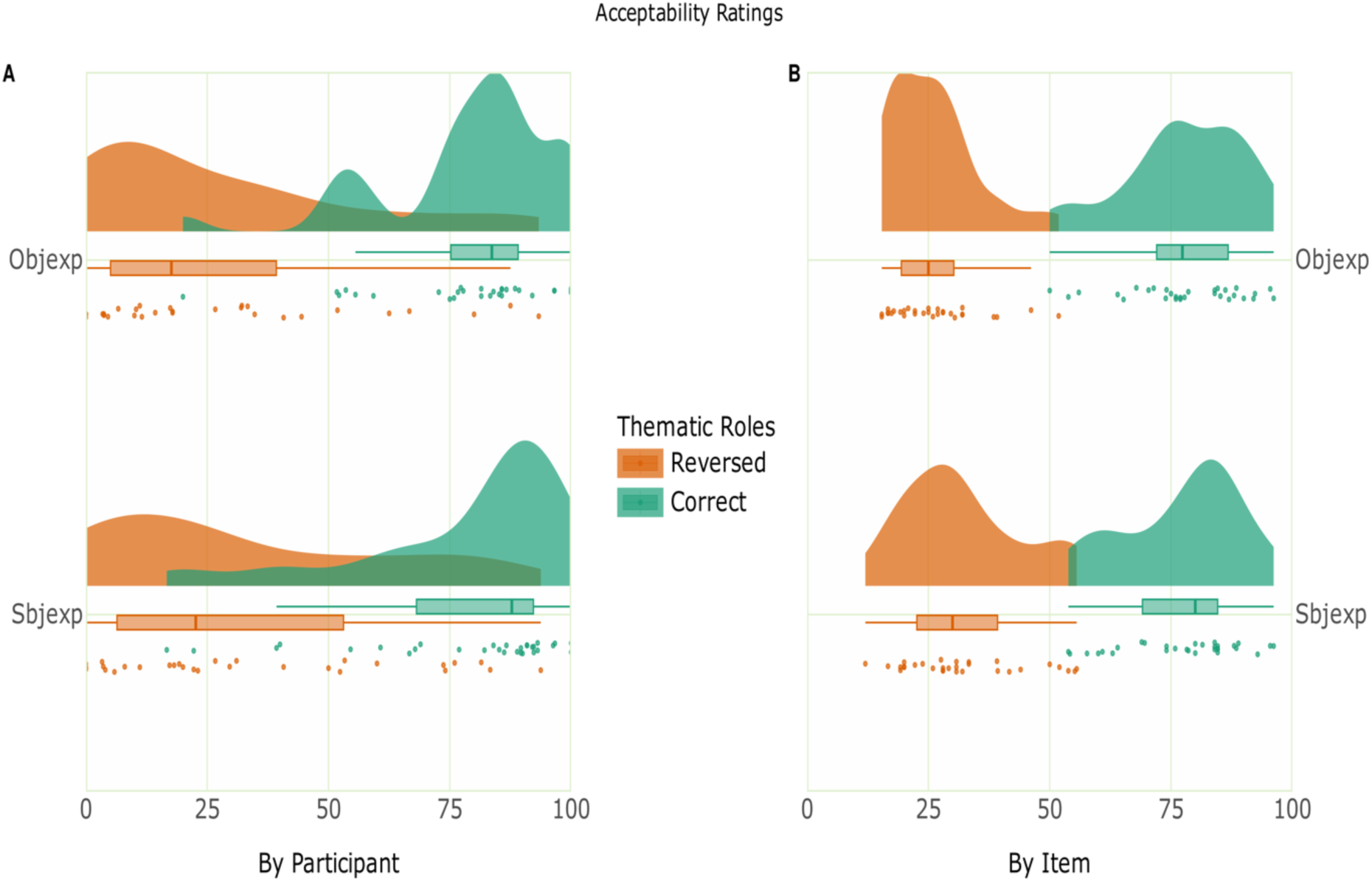
Raincloud plot of the acceptability ratings. Panel A shows the by-participant variability of acceptability ratings, with the individual data points representing the mean by-participant acceptability of each Verb type and Thematic roles combination. Panel B shows the by-item variability of acceptability ratings, with the individual data points representing the mean by-item acceptability of each Verb type and Thematic roles combination.

**Table 3:**
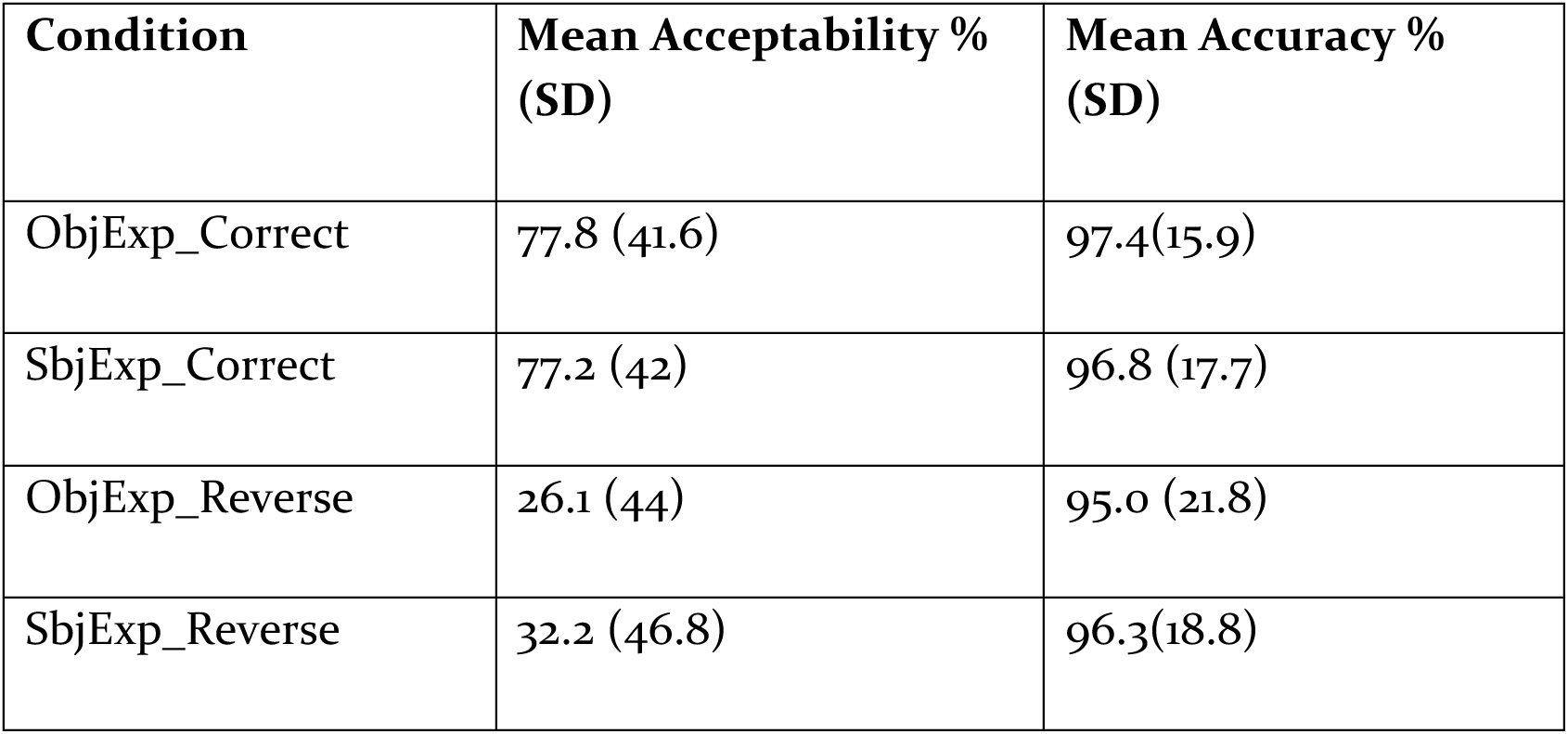
Mean acceptability ratings and probe detection accuracy.

| Condition | Mean Acceptability %<br>(SD) | Mean Accuracy %<br>(SD) |
| --- | --- | --- |
| ObjExp_Correct | 77.8 (41.6) | 97.4(15.9) |
| SbjExp_Correct | 77.2 (42) | 96.8 (17.7) |
| ObjExp_Reverse | 26.1 (44) | 95.0 (21.8) |
| SbjExp_Reverse | 32.2 (46.8) | 96.3(18.8) |

The behavioural acceptability and accuracy were analysed by fitting generalized linear mixed models using the lme4 package in R. Categorical fixed factors Centro sum contrasts (scaled sum contrasts for two-level factors). In the analysis of acceptability data, the statistical model included the fixed factors Verb type (SbjExp and ObjExp) and Thematic roles (Correct and Reverse), with random intercepts for participants and items, and by-participant random slopes for the effect of Verb type, Thematic roles and their interaction term. Type II Wald chi-square tests of the fitted model (AIC = 2789.21) of the acceptability data showed a main effect of Thematic roles (χ^2^(1) = 79.28, p < 0.001, s = 60.68).

In the analysis of probe detection accuracy, the statistical model included fixed effects of Verb type, Thematic roles, and their interaction, with random intercepts for participants and items, and by-participant random slopes for the effects of Verb type, Thematic roles, and their interaction. Type II Wald chi-square tests of the fitted model (AIC = 964.23) showed a main effect of Thematic roles (χ^2^(1) = 4.88, p = 0.02, s = 5.20).

### 4.2. ERP Data

The ERPs at the critical verb position are shown in Figure 2. Visual inspection of the ERP data showed that the role reversed conditions engendered a negativity effect followed by a positivity effect at the verb compared to their correct counterparts. In addition to considering ERP components that are relevant for reversal processing and their typical latencies based on previous literature, this qualitative observation based on visual inspection informed the choice of time-windows for analysis, namely 300–500 ms and 750-950. The single trial ERP mean amplitudes extracted in the analysis time-windows from a total of 3076 data epochs entered the analysis with a median of 769 trials per critical condition. The raw data collected during the experiment, the pre-processing pipeline used, and the pre-processed data are available in the data repository online. The ERPs at the sentence-initial noun and sentence-medial noun are not part of our hypotheses but are provided for reference in Figure S1 and Figure S2 in the supplementary material.

**Figure 2.**
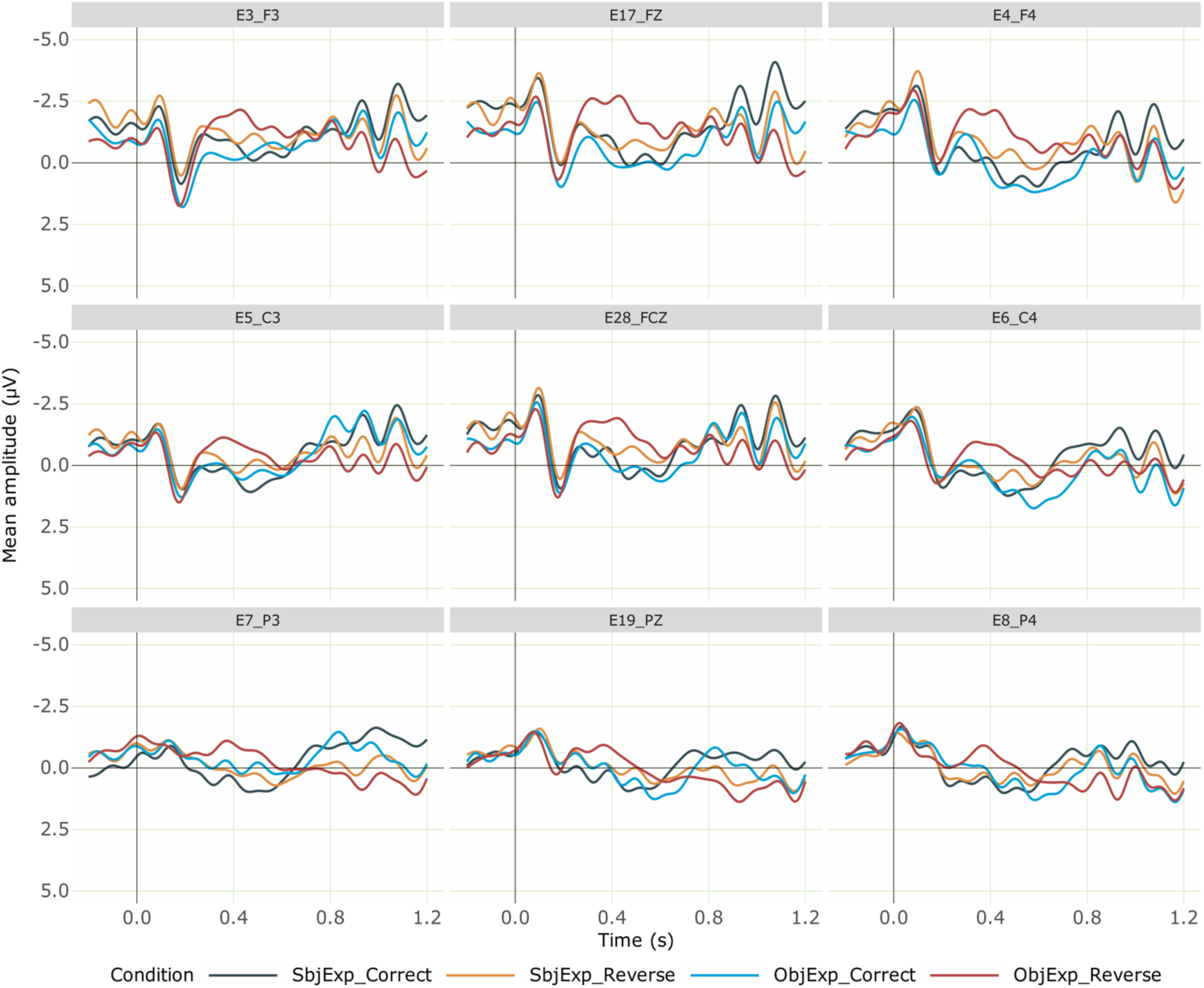
Grand averaged ERPs at the verb for the critical conditions from 30 participants. Negativity is plotted upwards; the time axis runs from −0.2 s to 1.2 s (i.e., - 200 ms to 1200 ms) with 0 being the onset of the critical verb. The dark blue line represents the correct SbjExp verb condition, while the orange line shows the role reversed SbjExp condition, which elicited a negativity followed by a late-positivity effect. The light blue line indicates the correct ObjExp verb condition and the red line represents the role reversed ObjExp verb condition, which elicited a negativity followed by a late-positivity effect.

#### 4.2.1. Planned Analysis

##### Time-window: 300-500 ms

We first computed a linear mixed-effects model m0 with the fixed factors Verb type, Thematic roles, ROI, the −200-0 ms pre-stimulus baseline mean amplitude as a covariate (scaled and centred), and by-participant and by-item random intercepts. The analysis code and full model outputs are available as R notebooks in the online analysis repository. Type II Wald Chi-squared tests on this model (AIC = 399178.29) showed main effects of Verb type (χ^2^(1) = 4.84, *p* = 0.02, s = 5.17) and Thematic roles (χ^2^(1) = 8.64, *p* = 0.003, s = 8.25), and an interaction of ROI x Thematic roles (χ^2^(1) = 13.15, p = 0.06, s = 3.86). Estimated marginal means on the response scale were computed using the emmeans package on the model to resolve the interaction. The pairwise contrasts of estimates for Thematic roles within each level of ROI revealed simple effects of Thematic role in the the left-frontal (estimate = −0.968, SE = 0.333, p = 0.003, s = 8.10), right-frontal (estimate = −1.143, SE = 0.333, p = 0.0006, s = 10.69), left-central (estimate = −0.672, SE = 0.333, p = 0.04, s = 4.52), right-central (estimate = −0.805, SE = 0.333, p = 0.01, s = 5.99), mid-parieto-occipital (estimate = −0.524, SE = 0.293, p = 0.07, s = 3.77) and mid-fronto-central (estimate = −1.195, SE = 0.333, p = 0.0003, s = 11.53) regions. The estimates for the thematic role reversed conditions were more negative than those for the correct thematic roles conditions in all the regions.

In order to verify if the model fit could be improved, we then computed another model m1 by making the random effects structure in m0 maximal, namely by including by-participant random slopes for the factor Verb, Thematic role and their interaction term. As mentioned earlier, a by-item random intercept specification is not identifiable for our data in view of how items were coded in the data, and therefore the by-item random intercept specification in m0 was retained as is in m1. Such a model would better account for within-participant ERP variance between conditions. Indeed, the model fit showed an improvement of (AIC = 398551.22), and type II Wald Chi-squared tests on this improved model showed a main effect of Thematic roles (χ^2^(1) = 4.28, p = 0.03, s = 4.69) and an interaction of ROI x Thematic roles (χ^2^(1) = 13.73, *p* = 0.05, s = 4.15). Estimated marginal means on the response scale were computed on the model to resolve the interaction, which revealed that the estimate for the role reversed condition was more negative in left-frontal (estimate = −0.938, SE = 0.404, p = 0.02, s = 5.63), right-frontal (estimate = −1.118, SE = 0.404, p = 0.005, s = 7.47), right-central (estimate = −0.774, SE = 0.403, p = 0.05, s = 4.18) and mid-fronto-central (estimate = −1.17, SE = 0.404, p = 0.003, s = 8.04) regions. The estimates for the thematic role reversed conditions were more negative than those for the correct thematic roles conditions in all the regions.

##### Time-window: 750-950 ms

Similarly to the analysis in the earlier time window, we first computed a linear mixed-effects model m0 in this time window with the fixed factors Verb type, Thematic roles, ROI, the −200-0 ms pre-stimulus baseline mean amplitude as a covariate (scaled and centred) and by-participant and by-item random intercepts. The type type II Wald Chi-squared tests on this model (AIC = 415046.03) showed an interaction of ROI x Thematic roles (χ^2^(1) = 14.228, p = 0.04, s = 4.40). The pairwise contrasts of estimates for Thematic roles within each level of ROI revealed simple effects of Thematic roles in left-parietal (estimate = 1.025, SE = 0.422, p = 0.01, s = 6.03) and mid-parieto-occipital (estimate = 0.855, SE = 0.380, p = 0.02, s = 5.35) regions. The estimates for the thematic role reversed conditions were more positive than those for the correct thematic roles conditions in all the regions.

In order to verify if the model fit could be improved, we computed another model m1 by making by-participant random slopes for the factors Verb type, Thematic roles and their interaction term. The type II Wald Chi-squared tests on this model (AIC = 414086) again showed a ROI x Thematic roles interaction (χ^2^(1) = 15.028, *p* = 0.03, s= 4.81). The pairwise contrasts of estimates for Thematic roles within each level of ROI revealed simple effects of Thematic roles in left-parietal (estimate = 1.134, SE =0.575, p = 0.04, s = 4.36) and mid-parieto-occipital (estimate = 0.964, SE = 0.545, p = 0.07, s = 3.69) regions. The estimates for the thematic role reversed conditions were more positive than those for the correct thematic roles conditions in all the regions.

In both time-windows, notwithstanding the subtle differences between the random intercepts and random slopes models, the overall pattern of results remains intact, namely that both the reversal conditions involving SbjExp verbs and ObjExp verbs elicited a biphasic effect: a negativity effect (300-500 ms) followed by a positivity effect (750-950 ms). Whilst a direct comparison of the ERPs of the two violation conditions with each other was not planned at the outset, the negativities evoked by the two violation conditions was of different magnitudes such that the difference between the effects was visually prominent, which we pursued further in a post-hoc analysis.

#### 4.2.2. Post-hoc analysis

Visual inspection of the data revealed that the reversal involving ObjExp verbs elicited a prominently larger negativity compared to the reversal involving SbjExp verbs. In view of this unplanned observation, we conducted a post-hoc analysis that addressed the question of whether there is some statistical evidence to support this visual difference in our data. Whilst an interaction of Verb type and Thematic roles was neither detected in the random intercepts model nor in the random slopes model of the ERP data in the 300-500 ms time-window, one crucial difference between the two models is that there was a main effect of Verb type in the random intercepts model, which was not detected in the random slopes model. The main effect of Verb type indeed suggests that the ERPs of the two verb types differ. However, in order to disentangle how they exactly differed, we computed estimated marginal means on the response scale using the emmeans package to resolve the Verb type x Thematic roles interaction in the model that detected both a main effect of Verb type and a main effect of Thematic roles, namely the random intercepts model m0 pertaining to the 300-500 ms time-window. Comparing the estimates of the two Verb types in each level of Thematic role revealed that the estimated mean amplitude for the ObjExp verbs in the reversal condition (estimate = - 1.224, SE = 0.370, p < 0.001, s = 10.06) was more negative than that for the SbjExp verbs in the reversal condition (estimate = −0.380, SE = 0.370, p = 0.30, s = 1.72). Similarly, the estimated means for the ObjExp verbs in the correct condition (estimate = −0.184, SE = 0.370, p = 0.61, s = 0.69) was more negative than that for the SbjExp verbs in the correct condition (estimate = 0.140, SE = 0.370, p = 0.70, s = 0.50). That is, the estimates for conditions with ObjExp verbs are generally more negative than those for conditions with SbjExp verbs in the 300-500 ms time-window. More interestingly for present purposes however, the contrast between the estimates of the two Verb types showed a simple effect of Verb type when Thematic roles were reversed (estimate = 0.843, SE = 0.363, p = 0.02, s = 5.63), whereas there was no such effect detected when Thematic roles were correct (estimate = 0.324, SE = 0.362, p = 0.37, s = 1.43). That is, there is converging statistical evidence in the data to support the visually prominent difference between the negativities evoked by the two violation conditions. In effect, this indicates that Verb type indeed appears to modulate the negativity effect evoked by reversal anomalies such that the effect is larger for ObjExp verbs than for SbjExp verbs in our data.

## 5. Discussion

We have presented an ERP experiment on Malayalam, which aimed to examine whether the verb type, namely SbjExp and ObjExp verbs, modulates the ERP correlates for thematic reversal anomalies in Malayalam. The behavioural task showed low acceptability ratings for the conditions in which the roles were anomalously reversed compared to their correct counterparts. Electrophysiological results for both SbjExp and ObjExp verbs in the thematic role reversed conditions revealed a biphasic N400–P600 pattern. Additionally, we observed a difference in the amplitude of the negativity effect, with ObjExp verbs eliciting a more pronounced negativity than SbjExp verbs. As reported above, previous research on reversal anomaly has reported different patterns: only N400, only P600 and N400-P600. As the present study reports a biphasic N400-P600 effect, in what follows, we discuss the functional interpretation of the biphasic N400-P600 effect that we found for the reversal conditions involving both SbjExp and ObjExp verbs.

### 5.1. N400-P600

As mentioned above, we observed a biphasic N400-P600 effect for TRAs involving SbjExp and ObjExp verbs in the present study. This finding aligns with several previous findings: German irresolvable reversal conditions with Act verbs (Schlesewsky and Bornkessel-Schlesewsky, 2009), reversal on Icelandic non-alternating verbs (Bornkessel-Schlesewsky et al., 2011) and English SbjExp verbs (Bourguignon et al., 2012). However, our results stand not only in contrast to findings from Chinese (Bornkessel-Schlesewsky et al., 2011) and English (Ehrenhofer et al., 2022), but also surprisingly from Turkish (Bornkessel-Schlesewsky et al., 2011), an overtly case-marked Altaic verb-final language with agglutinative morphology – features it shares with Malayalam. These studies reported only an N400 effect for reversal anomalies at the verb position. Bornkessel-Schlesewsky et al. (2011) interpreted the N400 effect found in the case of Turkish and Chinese based on word order dependence vs. independence for verb-argument linking. The absence of a P600 in these studies is said to be due to a structural ambiguity between a main clause versus a relative clause reading at the verb position, which arises from the lack of case marking on both arguments in the reversal condition in these studies. This ambiguity leads the processing system to perhaps opt for an alternative structural analysis, which in turn is said to make it a potentially resolvable anomaly (and thus no P600). As for the N400 effect found in English, Ehrenhofer et al. (2022) interpreted the effect as reflecting lexical predictability and integration of thematic role information into online sentence comprehension. In the following, we interpret the Malayalam finding in the light of the different accounts proposed in the above mentioned studies for an N400-P600 patternOur experimental design is particularly comparable to the German reversal condition in (2a) from Schlesewsky and Bornkessel-Schlesewsky (2009), in which TRA occurs at the sentence final verb after all arguments have already been processed, unlike in Icelandic in Bornkessel-Schlesewsky et al. (2011), where the critical verb position is sentence medial. Crucially, the arguments in (2a) are overtly case-marked, eliminating structural ambiguity and leaving no room for alternative interpretations. Similarly, in our study, both arguments are explicitly case marked, and the sentence structure does not allow for alternative analyses, further reinforcing the parallel with the German data.

3.

(a)…dass der Schalter den Techniker bedient that [the switch]:Nom [the technician]:Acc operates “…that the switch operates the technician.” (anomalous, for roles cannot be switched due to case marking information)
(b)…dass Schalter Techniker bedienen that switches technicians operate “…that technicians operate switches.”

(locally ambiguous, but not anomalous after reanalysis)

(German, Schlesewsky and Bornkessel-Schlesewsky, 2009)

Schlesewsky and Bornkessel-Schlesewsky (2009) interpreted the N400 effect within the framework of the eADM in terms of whether verb-argument linking depends on or is independent of word order. In languages like German, argument linking does not rely solely on word order; instead, it draws on a range of sequence-independent cues, including case marking and animacy. A processing conflict arises during the compute linking step due to mismatches between case, word order, and animacy information. For example, in sentence (2a), the nominative case of NP1 suggests that *the switch* should be interpreted as the actor, whereas animacy cues favour interpreting *the technician* as the actor. Similarly, in sentence (2b), word order supports an actor interpretation of *the switches* (due to underspecified case marking, the plural noun can be a subject or object), while animacy again favours *the technicians.* Such a conflict engenders an N400 effect.

While the interpretation proposed for German studies partially accounts for our findings, it does not fully explain the entire pattern of results. Specifically, the N400 interpretation they provided fits well with our reversal condition involving SbjExp verbs but not for the reversal condition involving ObjExp verbs. In terms of the interpretation provided by Bornkessel-Schlesewsky et al. (2011) based on the eADM, in the reversal condition with SbjExp verb (refer back to Table 2), the nominative case of NP1 together with the fact that the argument is sentence-initial call for an actor interpretation for NP1, while the animacy information (inanimate) is more compatible with an undergoer interpretation for NP1. Similarly for the animate argument at NP2, case information calls for an undergoer reading and animacy information is more compatible with an actor reading. Hence a processing problem already occurs in the Compute Linking step due the conflict between animacy and case information, thereby triggering the observed N400 effect. In contrast, the reversal condition with ObjExp verbs does not present a conflict between case marking and animacy cues. In these sentences, the nominative case on the first, animate argument supports an Actor role, and animacy cues are consistent with this interpretation. The second argument is inanimate and marked with accusative case, aligning with an Undergoer role. As a result, no conflict arises during the Compute Linking stage in terms of the eADM, unlike in SbjExp verb reversals. Thus, only a P600 effect reflecting difficulties at the generalized mapping stage would be expected, without a preceding N400. However, we not only observed an N400 effect for ObjExp verbs, but rather the effect was larger for these verbs compared to SbjExp verbs. Bourguignon et al. (2012) offer an updated account of TRAs within the Eadm framework, arguing that the presence or absence of the N400 also depends on the extent to which lexical-level processing is engaged during sentence comprehension. They reported a biphasic N400–P600 pattern in response to reversal conditions involving SbjExp verbs, in contrast to Act verbs. In our study, we observed a similar biphasic pattern for both SbjExp and ObjExp verbs. Following the eADM-based interpretation of Bourguignon et al. (2012), we can also interpret the N400 effect in our data as an index of thematic reanalysis process. Specifically, the early assignment of agent/patient roles is revised as soon as more fine-grained thematic information based on the verb’s logical-structure is accessed during the argument linking step. In the reversal conditions, the initially assigned actor-subject and patient-object roles must be reinterpreted as experiencer-subject and experiencer-object, respectively. The higher amplitude N400 observed for reversal conditions involving ObjExp verbs, as compared to SbjExp verbs, suggests that reanalysing a patient-object as an experiencer is potentially more effortful than reanalysing an agent-subject as an experiencer. This asymmetry can be attributed to the thematic properties associated with these roles: in SbjExp constructions, the experiencer retains many prototypical agent-like features, whereas in ObjExp constructions, it is the stimulus or theme that assumes more agent-like characteristics (Dowty, 1991). As a result, shifting from an agent interpretation to an experiencer interpretation is cognitively less demanding than shifting from a patient interpretation to an experiencer interpretation. The subsequent P600 effect is taken to reflect a binary categorization process, distinguishing between anomalous and non-anomalous conditions. However, this interpretation becomes problematic, as discussed in the “Electrophysiology of Thematic Reversal Anomalies” section, when we consider data from Kyriaki et al. (2020). Kyriaki et al. (2020), when replicating the study by Bourguignon et al. (2012), observed an N400 effect even for Act verbs. The presence of an N400 in the action verb condition challenges the reanalysis-based interpretation, as no thematic reanalysis should be required in these cases. Specifically, the initially computed actor role remains valid (albeit assigned to the wrong event participant) upon accessing the verb’s thematic structure, suggesting that reanalysis is not necessary for Act verbs in reversal conditions. Kyriaki et al. (2020) employed multiple modalities (visual and auditory), tasks (judgment and comprehension) and analyses methods (ANOVA/MEM) which introduced substantial variability. As a result, the findings are difficult to interpret clearly. However, a recent study by Ehrenhofer et al. (2022) on English also found an N400 for reversal conditions in both verb medial (NP-V-NP) and verb final positions (NP-NP-V). These findings suggest that the interpretation of the N400 as reflecting a reanalysis of previously computed thematic roles may need to be reconsidered.

Another possibility for explaining the N400-P600 found in our study is the account provided by Delogu et al. (2025) based on the retrieval-integration framework of Brouwer et al. (2012). They interpreted the biphasic N400-P600 pattern found for linguistic anomalies as reflecting increased processing effort during both retrieval and integration stages. According to the account, the N400 is modulated by factors such as lexical association and contextual expectancy that determine the ease of accessing word meaning from long-term memory. The P600 component in this account is an index of increased processing demands during the integration of the activated lexical information into the existing current mental representation of an unfolding sentence. Delogu et al. (2025) and Brouwer & Crocker (2017) suggested that spatio-temporal overlap between components can mask or attenuate the P600, especially when retrieval and integration happen simultaneously, but also due to task-induced depth of processing and the nature of the anomaly. The P600 effect observed in our study for both reversal conditions involving SbjExp and ObjExp verbs can be interpreted as a marker of integration difficulty. This interpretation is further supported by the fact that our study used an acceptability judgment task, which likely encouraged deeper processing of sentence meaning. However, the interpretation of the N400 offered by Delogu et al. (2025) and Brouwer et al. (2012) does not fully account for the N400 effect observed in our study nor those found in some of the other studies such as Bornkessel-Schlesewsky et al. (2011), Bourguignon et al. (2012) and Kyriaki et al. (2020). The arguments used in our stimuli were semantically and conceptually compatible with the verbs, indicating a high degree of lexical association. Under retrieval-integration view, an N400 effect would not be expected. Yet, we observed a robust N400 effect, suggesting that additional factors beyond simple lexical association may be contributing to the observed processing difficulty.

In sum, the accounts discussed above for interpreting N400s found in the context of TRAs lead to contradictory predictions, and none of them provides a parsimonious account for the range of N400 effects reported across languages for TRAs. In this regard, the neurobiologically plausible model of language-related negativities proposed by Bornkessel-Schlesewsky & Schlesewsky (2019) offers a promising approach to reconciling the seemingly contradictory N400 effects observed for TRAs across languages. This model interprets all language-related negativities based on the predictive coding framework, whereby the brain continuously generates predictions about upcoming linguistic input. The amplitude modulations of all negativities including the N400 are said to reflect the precision-weighted prediction error signals, i.e., prediction errors weighted by the relevance of the information source leading to the error. According to this account, precision is closely tied to cue validity within a given language. This means that the more valid or reliable a linguistic cue is for interpreting sentence structure or meaning, the more it influences how strongly the brain updates its internal predictive model during comprehension. Drawing on the predictive coding framework, the authors account for cross-linguistic differences in the processing of TRAs. They argue that although TRAs reliably generate prediction errors, these are not always reflected in an N400 amplitude modulation. Instead, they propose that the N400 amplitude modulation reflects updation of the internal model rather than merely signalling the presence of a prediction error. A key factor in whether an error leads to model updating is the precision of the error signal, which is influenced by the cue validity of the mismatch-inducing feature (e.g., animacy, word order, case marking, etc). If the prediction error arises from a cue that is highly relevant for sentence interpretation in a given language, the parser is more likely to update its internal model. By contrast, if the error stems from a less relevant cue, the parser may detect the mismatch but not treat it as significant enough to trigger model updating. Further, with converging evidence from Bornkessel-Schlesewsky et al. (2011) and Bourguignon et al. (2012), Bornkessel-Schlesewsky & Schlesewsky (2019) argued that the relevance of a cue for sentence interpretation may change depending on the circumstances in which it is encountered. Findings from Kyriaki et al. (2020) provide converging evidence that cue relevance is not fixed but can vary depending on factors such as task demands (e.g., acceptability judgment vs. comprehension tasks) and modality (visual vs. auditory presentation). Similarly, findings from Ehrenhofer et al. (2022) demonstrate that even subtle differences in stimulus materials can significantly affect whether prediction-related cues – such as argument role information – are effectively used during sentence processing.

In light of this, the N400 effect observed in our study can be interpreted as reflecting the updating of the internal generative model. Specifically, N400 amplitude indicates whether the parser actually updates its internal generative model of language in response to a prediction error (Bornkessel-Schlesewsky & Schlesewsky, 2019). The preverbal arguments in our stimuli allow for multiple plausible continuations, including simple and complex transitive constructions with an Act verb or experiencer verb, or even relative clauses (refer sentences 3a to 3d). The case marking and animacy information of the two arguments are highly valid and reliable cues in Malayalam allowing for a more precise prediction of the verb. The presence of a SbjExp verb following an inanimate subject and animate object (SbjExp_Reverse), and the presence of an ObjExp verb following an animate subject and inanimate object (ObjExp_Reverse) induces a conflict with prior predictions about the verb, and results in a prediction error. Once prediction error is encountered, the parser tries to update the internal model depending on how precise the prediction error is, in order to generate a revised set of predictions to accommodate the unexpected input.

4.

(a) Maranam Indiraye keezhadakki. Death (In.Nom) Indira(An.F)-Acc conquer-Pst “Death conquered Indira.”
(b) Maranam Indiraye thediyethi. Death (In.Nom) Indira(An.F)-Acc search-come-Pst “Death came in search of Indira.”
(c) Subeena ulsavathe avaganichu. Subeena(An.F.Nom) festival (In)-Acc ignore-Pst “ Subeena ignored the festival.”
(d) Subeena ulsavathe cholli santhoshichu. Subeena(An.F.Nom) festival (In)-Acc recite be.happy-Pst “Subeena rejoiced because of the festival.”

According to this account, the amplitude difference in N400 found between TRAs involving SbjExp and ObjExp verbs can be interpreted as reflecting differences in the precision of prediction error^2^. In ObjExp_Reverse sentences, their Animate-Nom-NP1—Inanimate-Acc-NP2 structure permits a limited set of plausible continuations, most notably a SbjExp verb that would enable interpreting the animate nominate NP1 as an experiencer and the inanimate accusative NP2 as the stimulus (refer to sentences 3c and 3d). This expectation arises because inanimate accusative objects are limited to very specific exceptions of differential object marking in Malayalam (Aissen, 2003; De swart, 2007; Egger, 2016; García García et al., 2018), thereby strongly constraining the range of verbs that can plausibly follow this configuration. Thus, the convergence of animacy, case and positional cues make the initial prediction for a specific kind of verb relatively more precise. When the clause final verb instead turns out to be an ObjExp verb requiring inverse linking, this expectation must be revised, forcing a reweighting of the cues used for argument interpretation. Consequently, this shift in cue relevance amplifies the precision-weighted prediction error at the verb, which necessitates updating the internal generative model and evokes a larger N400 effect for TRAs involving ObjExp verbs. In contrast, in the case of SbjExp_Reverse sentences, their Inanimate-Nom-NP1—Animate-Acc-NP2 structure permits a wider range of plausible constructions (refer to sentences 3a and 3b). Thus, the prediction for a specific kind of verb would be less precise, thus resulting in a relatively smaller prediction error, in turn resulting in a smaller N400 effect. Such an interpretation would further be in line with Ehrenhofer et al. (2022), who argued that the presence or absence of an N400 effect in the context of TRAs appears to depend on the strength and precision of predictions about the upcoming verb. In effect, our findings suggest that cue relevance for argument interpretation is modulated by the linking properties associated with the verb class. In the case of ObjExp verbs, their inverse linking nature alters the predicted mapping between grammatical functions and thematic roles, shifting the focus from canonical prominence-based cues (such as animacy and case of the arguments) to verb-specific linking information. The observed larger N400 amplitude difference therefore reflects the system adjusting cue weighting when the anticipated form-to-meaning mapping is overridden by the inverse linking requirements of the ObjExp verbs. Our findings thus provide converging support for the observation made by Bornkessel-Schlesewsky & Schlesewsky (2019) that the relevance of a cue for sentence interpretation may change depending on the circumstances in which it is encountered.

The P600 observed in our study can be accounted for by the various accounts outlined above: it may reflect the irresolvability of the conflict (Schlesewsky and Bornkessel-Schlesewsky, 2009), integration difficulty (Brouwer et al., 2012; Brouwer & Crocker, 2017; Delogu et al., 2025; Zhang et al., 2025), or it may be interpreted as a categorization-related P600 (Bornkessel-Schlesewsky et al., 2011). The P600 can also be explained by other accounts, especially as a marker of monitoring process, encompassing both the conflict and reprocessing (Meerendonk et al., 2013; Vissers et al., 2013), repair and reanalysis process (Kos et al., 2010; Puhacheuskaya, 2021), or a task dependent P600 (Brouwer & Crocker, 2017; Sassenhagen et al., 2014). However, the interpretation most consistent with our N400 interpretation would be considering the P600 as a reflection of the behavioural consequences of predictive coding response (Bornkessel-Schlesewsky & Schlesewsky, 2019). After detecting a prediction error, the parser initiates an update, in which it down-weights the prior prediction, recalibrates its cue weighting, and generates a revised set of predictions to accommodate the unexpected input. However, even after model updating, if the parser no longer yields a coherent interpretation, the sentence is categorized as anomalous, which is reflected in P600 effect. The acceptability judgment task we used may have also accelerated this process.

## 6. Conclusion

The present study explored the processing of TRAs involving SbjExp and ObjExp verbs in Malayalam. The results revealed a biphasic N400-P600 effect for both thematic role reversed conditions involving SbjExp and ObjExp verbs, suggesting that despite their syntactic and semantic differences, they are processed qualitatively similarly when a thematic reversal anomaly is involved. However, we also found an N400 amplitude difference between the verb types in the post-hoc analysis, whereby TRAs involving ObjExp verbs evoked a larger N400 effect in comparison to TRAs involving SbjExp verbs. We suggest that this quantitative difference observed for ObjExp verbs is due to the inverse linking of grammatical function and thematic roles associated with these verbs. In other words, the parser recalibrates cue weighting when the expected form-to-meaning mappings are overridden by the inverse linking properties of object experiencer verbs. These findings therefore suggest that verb-specific linking properties shape cue relevance during argument interpretation, highlighting how fine-grained differences within subtypes of experiencer verbs modulate the neurocognitive processes underlying sentence comprehension.

## Data availability statement

The statistical analysis code and full model outputs of all the analyses reported are available as R notebooks here: https://doi.org/10.5281/zenodo.19283265

## Ethics statement

The studies involving humans were approved by Institute Ethical Committee of the Indian Institute of Technology Ropar. The studies were conducted in accordance with the local legislation and institutional requirements. The participants provided their written informed consent to participate in this study.

## Author contributions

**SS**: Resources, Conceptualization, Writing original draft, Project administration, Validation, Formal analysis, Investigation, Methodology, Software, Writing-review & editing, Data curation, Visualization. **RM**: Supervision, Methodology, Funding acquisition, Formal analysis, Writing - review & editing, Software, Visualization, Data curation, Validation, Conceptualization. **IBS**: Methodology, Formal Analysis, Writing – review & editing **MS:** Methodology, Formal Analysis, Writing – review & editing **KKC**: Conceptualization, Methodology, Funding acquisition, Validation, Project administration, Supervision, Writing – review & editing.

## Footnotes

1 We did not include Act verbs in our design, due to potential confounds they may introduce in view of differential object marking in Malayalam syntax (de Swart, 2003; 2007; Asher, 2013). For instance, inanimate object arguments need to be marked accusative in experiencer verbs constructions, (as in “John karine(car-Acc) snehichu. – John loved the car”), but by contrast, marking an inanimate object argument accusative for Act verbs would be ungrammatical in Malayalam; instead inanimate objects have to be in the unmarked (nominative) form for Act verbs (as in “John karu odichchu” – “John drove the car”).

2 The larger N400 amplitude observed for TRAs involving ObjExp is reminiscent of findings from Bornkessel-Schlesewsky et al. (2011) for Turkish. In that study, reversal sentences involving verbs that require an inanimate Actor and an animate undergoer elicited a larger N400 effect.

## Notes

https://doi.org/10.5281/zenodo.19283265

### Competing Interest Statement

The authors have declared no competing interest.

