## Supplementary Material for "Verb-Specific Linking Properties Modulate the N400 Effect: Evidence from Thematic Reversal Anomalies in Malayalam"

### A1: Critical sentences used in the study

| **SbjExp_Correct_C** | **#SbjExp_Reverse_X** |
|---|---|
| അമീർ സമ്മാനത്തെ സ്നേഹിച്ചു.<br>Ameer sammanthe snehichu.<br>Ameer(Nom) gift-Acc love-Pst<br>Ameer loved the gift. | സമ്മാനം അമീറിനെ സ്നേഹിച്ചു.<br>Sammanam Ameerine snehichu.<br>Gift(Nom) Ameer-Acc love-Pst<br>Gift loved Ameer |
| അജിത അവകാശത്തെ സ്നേഹിച്ചു.<br>Ajitha avakashathe snehichu.<br>Ajitha(Nom) rights-Acc love-Pst<br>Ajitha loved rights | അവകാശം അജിതയെ സ്നേഹിച്ചു.<br>Avakasham Ajithaye snehichu.<br>Rights(Nom) Ajtha-Acc love-Pst<br>Rights loved Ajitha. |
| ഗോവിന്ദ് അവകാശത്തെ വെറുത്തു.<br>Govindh avakshathe veruthu.<br>Govindh(Nom) rights-Acc hate-Pst<br>Govindh hated rights | അവകാശം ഗോവിന്ദിനെ വെറുത്തു.<br>Avakasham Govindhine veruthu.<br>Rights(Nom) Govindh-Acc hate-Pst<br>Rights hated Govindh |
| ഗ്രീഷ്മ സമ്മാനത്തെ വെറുത്തു.<br>Greeshma sammanthe veruthu.<br>Greeshma(Nom) gift-Acc hate-Pst<br>Greeshma hated gift. | സമ്മാനം ഗ്രീഷ്മയെ വെറുത്തു.<br>Sammanam Greeshmaye veruthu.<br>Gift(Nom) Greeshma-Acc hate-Pst<br>Gift hated Greeshma |
| ഗൗതം അപകടത്തെ പുച്ഛിച്ചു.<br>Gautham apakadathe puchichu.<br>Gautham(Nom) danger-Acc scorn-Pst<br>Gautham scorned danger | അപകടം ഗൗതത്തെ പുച്ഛിച്ചു.<br>Apakadam Gauthathe puchichu.<br>Danger(Nom) Gautham-Acc scorn-Pst<br>Danger scorned Gautham. |
| ഗായത്രി പദവിയെ പുച്ഛിച്ചു.<br>Gayathri padhaviye puchichu<br>Gayathri(Nom) position-Acc scorn-Pst<br>Gayathri scorned at position | പദവി ഗായത്രിയെ പുച്ഛിച്ചു.<br>Padhavi Gayathriye puchichu.<br>Position(Nom) Gayathri-Acc scorn-Pst<br>Position scorned at Gayathri. |
| സഹീർ പദവിയെ ബഹുമാനിച്ചു.<br>Saheer padhaviye bahumanichu. | പദവി സഹീറിനെ ബഹുമാനിച്ചു.<br>Padhavi Saheerine bahumanichu. |

| | |
|---|---|
| Saheer(Nom) position-Acc respect-Pst<br>Gayathri respected posiiton | Position(Nom) Saheer-Acc respect-Pst<br>Position respected Gayathri |
| രേണുക അന്വേഷണത്തെ ബഹുമാനിച്ചു.<br>Renuka anweshanthe bahumanichu.<br>Renuka(Nom) investigation-Acc respect-Pst<br>Renuka respected the investigation | അന്വേഷണം രേണുകയെ ബഹുമാനിച്ചു.<br>Anweshanam Renukaye bahumanichu.<br>Investigation(Nom) Renuka-Acc respect-Pst<br>Investigation respected Renuka |
| കുമാർ അപകടത്തെ നിന്ദിച്ചു}<br>Kumar apakadathe nindhichu.<br>Kumar(Nom) danger-Acc denounce-Pst<br>Kumar denounced the danger. | അപകടം കുമാറിനെ നിന്ദിച്ചു.<br>Apakadam Kumarine nindhichu.<br>Danger(Nom) Kumar-Acc denounce-Pst<br>Danger denounced Kumar. |
| ഇന്ദിര അന്വേഷണത്തെ നിന്ദിച്ചു.<br>Indira anweshanam nindhichu.<br>Indira(Nom) investigation-Acc denounce-Pst<br>Indira denounced investigation | അന്വേഷണം ഇന്ദിരയെ നിന്ദിച്ചു<br>Anweshanam Indhiraye nindhichu.<br>Investigation(Nom) Indira-Acc denounce-Pst<br>Investigation denounced Indira. |
| ഉസ്മാൻ ഇരുട്ടിനെ ആദരിച്ചു.<br>Usman iruttine aadharichu.<br>Usman(Nom) darkness-Acc respect-Pst<br>Usman respected darkness. | ഇരുട്ട് ഉസ്മാനെ ആദരിച്ചു<br>Irutt Usmane aadharichu.<br>Darkness(Nom) Usman-Acc respect-Pst<br>Darkness respected Usman. |
| ഐശ്വര്യ സംഗീതത്തെ ആദരിച്ചു<br>Aishwarya sangeethahe aadharichu.<br>Aishwarya(Nom) music-Acc respect-Pst<br>Aishwarya respected music | സംഗീതം ഐശ്വര്യയെ ആദരിച്ചു<br>Sangeetham Aishwaryaye aadharichu<br>Music(Nom) Aishwarya-Acc respect-Pst<br>Music respected Aishwarya |
| കണ്ണൻ ഇരുട്ടിനെ വിസ്മരിച്ചു<br>Kannan iruttine vismarichu.<br>Kannan(Nom) darkness-Acc forget-Pst<br>Kannan forgot the darkness | ഇരുട്ട് കണ്ണനെ വിസ്മരിച്ചു.<br>Irutt Kannane vismarichu.<br>Darkness(Nom) Kannan-Acc forget-Pst<br>Darkness forgot Kannan |
| കാവേരി സംഗീതത്തെ വിസ്മരിച്ചു.<br>Kaveri sangeethathe vismarichu.<br>Kaveri(Nom) music-Acc forget-Pst<br>Kaveri forgot the music. | സംഗീതം കാവേരിയെ വിസ്മരിച്ചു.<br>Sangeetham Kaveriye vismarichu.<br>Music(Nom) Kaveri-Acc forget-Pst.<br>Music forgot Kaveri. |

| | |
|---|---|
| ആദർഷ് ജീവിതത്തെ പ്രണയിച്ചു.<br>Adharsh jeevithathe prananyichu.<br>Adharsh(Nom) life-Acc love-Pst<br>Adarsh loved life | ജീവിതം ആദർഷിനെ പ്രണയിച്ചു<br>Jeevitham Adharshine pranayichu.<br>Life(Nom) Adharsh-Acc love-Pst<br>Life loved Adharsh. |
| ഇന്ദിര മരണത്തെ പ്രണയിച്ചു<br>Indira maranthe pranayichu.<br>Indira(Nom) death-Acc love.Pst<br>Death loved Indira . | മരണം ഇന്ദിരയെ പ്രണയിച്ചു<br>Maranam Indiraye pranayichu.<br>Death(Nom) Indira-Acc love-Pst<br>Death loved Indira. |
| കലേഷ് യാത്രയെ പ്രേമിച്ചു<br>Kalesh yaathraye pranayichu.<br>Kalesh(Nom) travel-Acc love-Pst<br>Kalesh loved travel. | യാത്ര കലേഷിനെ പ്രേമിച്ചു}<br>Yathra Kaleshine pranayichu.<br>Travel(Nom) Kalesh-Acc love-Pst<br>Travel loved Kalesh. |
| കല്യാണി മരണത്തെ പ്രേമിച്ചു<br>Kalyani maranthe premichu.<br>Kalyani(Nom) death-Acc love.Pst<br>Kalyani loved darkness | മരണം കല്യാണിയെ പ്രേമിച്ചു<br>Maranam Kalyaniye premichu.<br>Death(Nom) Kalyani-Acc love-Pst<br>Darkness loved Kalyani |
| ആഷിഖ് ജീവിതത്തെ ആശിച്ചു<br>Aashique jeevithathe aashichu.<br>Aashique(Nom) life-Acc desire-Pst<br>Aashique desired life. | ജീവിതം ആഷിഖിനെ ആശിച്ചു.<br>Jeevitham Aashiqine aashichu.<br>Life(Nom) Aashique-Acc desire-Pst<br>Life desired Aashique. |
| ആരതി യാത്രയെ ആശിച്ചു<br>Aarathy yaathraye aashichu.<br>Aarathy(Nom) travel-Acc desire-P<br>Aarathy desired travel | യാത്ര ആരതിയെ ആശിച്ചു.<br>Yaathra aarathiye aashichu.<br>Travel(Nom) Aarathy-Acc desire-Pst<br>Travel desired Aarathy |
| അനില കളരിയെ സ്നേഹിച്ചു<br>Anila kalariye snehichu.<br>Anila(Nom) Kalari-Acc love-Pst<br>Anila loved Kalari | കളരി അനിലയെ സ്നേഹിച്ചു.<br>Kalari Anilaye snehichu.<br>Kalari(Nom) Anila-Acc love-Pst<br>Kalari loved Anila. |
| രൂപേഷ് കളരിയെ പ്രണയിച്ചു<br>Roopesh kalariye pranayichu. | കളരി രൂപേഷിനെ പ്രണയിച്ചു<br>Kalari Roopeshine pranayichu. |

| | |
|---|---|
| Roopesh(Nom) Kalari-Acc love-Pst<br>Roopesh loved Kalari. | Kalari(Nom) Roopesh-Acc love-Pst<br>Kalari loved Roopesh. |
| വിജയ് നൃത്തത്തെ വിസ്മരിച്ചു<br>Vijay nrithathe vismarichu.<br>Vijay(Nom) dance-Acc forget-Pst<br>Vijay forgot dance. | നൃത്തം വിജയിയെ വിസ്മരിച്ചു<br>Nritham Vijayiye vismarichu.<br>Dance(Nom) Vijay-Acc forget-Pst<br>Dance forgot Vijay. |
| നിമിഷ നൃത്തത്തെ ആശിച്ചു<br>Nimisha nrithathe aashichu.<br>Nimisha(Nom) dance-Acc desire-Pst<br>Nimisha desired dance. | നൃത്തം നിമിഷയെ ആശിച്ചു<br>Nritham Nimishaye aashichu.<br>Dance(Nom) Nimisha-Acc desire-Pst<br>Dance desired Nimisha. |
| റുബീന ചെണ്ടയെ പ്രേമിച്ചു<br>Rubeena chendaye premichu.<br>Rubeena(Nom) drums-Acc love-Pst<br>Rubeena loved drums. | ചെണ്ട റുബീനയെ പ്രേമിച്ചു<br>Chenda Rubeenaye premichu.<br>Drums(Nom) Rubeena-Acc love-Pst<br>Drums loved Rubeena. |
| ജീവൻ ചെണ്ടയെ ബഹുമാനിച്ചു.<br>Jeevan chendaye bahunanichu.<br>Jeevan(Nom) drums-Acc respect-Pst<br>Jeevan respected drums. | ചെണ്ട ജീവനെ ബഹുമാനിച്ചു<br>Chenda Jeevane bahumanichu.<br>Drums(Nom) Jeevan-Acc respect-Pst<br>Drums respected Jeevan. |
| മനോജ് ഉത്സവത്തെ ആദരിച്ചു<br>Manoj utsavathe aadharichu.<br>Manoj(Nom) celebration-Acc respect-Pst<br>Manoj respected celebration | ഉത്സവം മനോജിനെ ആദരിച്ചു<br>Utsavam Manojine aadharichu.<br>Celebration(Nom) Manoj-Acc respect-Pst<br>Celebration respected Manoj |
| സുബീന ഉത്സവത്തെ പുച്ഛിച്ചു<br>Subeena Utsavathe puchichu.<br>Subeena(Nom) celebration-Acc scorn-Pst<br>Subeena scorned celebration | ഉത്സവം സുബീനയെ പുച്ഛിച്ചു.<br>Utsavam Subeenaye puchichu.<br>Celebration(Nom) Subeena-Acc scorn-Pst<br>Celebration scorned Subeena. |
| ജ്യോതിഷ് വ്യായാമത്തെ നിന്ദിച്ചു<br>Jyothish vyayamathe nindhichu.<br>Jyothish(Nom) exercise-Acc denounce-Pst<br>Jyothish denounced exercise | വ്യായാമം ജ്യോതിഷിനെ നിന്ദിച്ചു.<br>Vyayam Jyothishine nindhichu<br>Exercise(Nom) Jyothish-Acc denounce-Pst<br>Exercise denounced Jyothish |

| | |
|---|---|
| ഗ്രീഷ്മ വ്യായാമത്തെ വെറുത്തു<br>Greeshma Vyayamathe veruthu.<br>Greeshma(Nom) exercise-Acc hate-Pst<br>Greeshma hated exercise | വ്യായാമം ജ്യോതിഷിനെ വെറുത്തു<br>Vyayam Jyothishine veruthu.<br>Exercise(Nom) Jyothish hate-Pst<br>Exercise hated Greeshma |

| **ObjExp_Correct_C** | **#ObjExp_Reverse_X** |
|---|---|
| സമ്മാനം അമീറിനെ ആഹ്ലാദിപ്പിച്ചു .<br>Sammanam Ameerine ahladhippichu<br>Gift(Nom) Ameer-Acc please-Pst<br>The gift pleased Ameer. | അമീർ സമ്മാനത്തെ ആഹ്ലാദിപ്പിച്ചു.<br>Ameer sanmmanthe ahladhippichu.<br>Ameer(Nom) gift-Acc please.Pst<br>The Ammer plesed gift |
| അവകാശം അജിതയെ ആഹ്ലാദിപ്പിച്ചു.<br>Avakasham Ajithaye ahaldhippichu.<br>Rights(Nom) Ajitha-Acc please-Pst<br>Rights pleased Ajitha | അജിത അവകാശത്തെ ആഹ്ലാദിപ്പിച്ചു.<br>Ajitha avakashathe ahladhippichu.<br>Ajitha(Nom) rights-Acc please-Pst<br>Ajitha pleased rights |
| അവകാശം ഗോവിന്ദിനെ രസിപ്പിച്ചു<br>Avakasham Govindhine rasippichu.<br>Rights(Nom) Govindh-Acc please-Pst<br>Rights pleased Govindh. | ഗോവിന്ദ് അവകാശത്തെ രസിപ്പിച്ചു<br>Govindh avakashathe rasippichu<br>Govindh(Nom) rights-Acc please-Pst<br>The freedom delighted Govindh. |
| സമ്മാനം ഗ്രീഷ്മയെ രസിപ്പിച്ചു.<br>Sammanam Greeshmaye rasippichu.<br>Gifts(Nom) Greeshma-Acc please-Pst<br>Gift pleased Greeshma | ഗ്രീഷ്മ സമ്മാനത്തെ രസിപ്പിച്ചു .<br>Greeshma sammanthe rasippichu.<br>Greeshma(Nom) gifts-Acc please-Pst<br>Gift pleased Greeshma |
| അപകടം ഗൗതത്തെ മുഷിപ്പിച്ചു<br>Apakadam Gauthathe mushippichu.<br>Danger(Nom) Gautham-Acc upset-Pst<br>The danger disturbed Gautham. | ഗൗതം അപകടത്തെ മുഷിപ്പിച്ചു<br>Gautham apakadathe mushippichu.<br>Gautham(Nom) danger-Acc upset-Pst<br>The danger disturbed Gautham. |
| പദവി ഗായത്രിയെ മുഷിപ്പിച്ചു<br>Padhavi Gayathriye mushippichu. | ഗായത്രി പദവിയെ മുഷിപ്പിച്ചു<br>Gayathri padhaviye mushippichu |

| | |
|---|---|
| Position(Nom) Gayathri-Acc upset-Pst<br>Position upset Gayathri | Gayathri(Nom) position-Acc upset-Pst<br>Gayathri upset position. |
| പദവി സഹീറിനെ ആനന്ദിപ്പിച്ചു<br>Padhavi Saheerine anadhippichu<br>Position(Nom) Saheer-Acc please-Pst<br>Position pleased Saheer. | സഹീർ പദവിയെ ആനന്ദിപ്പിച്ചു<br>Saheer padhaviye anandhippichu<br>Saheer(Nom) padhavi-Acc please-Pst<br>Saheer pleased position. |
| അന്വേഷണം രേണുകയെ ആനന്ദിപ്പിച്ചു<br>Anweshanam renukaye aanadhippichu.<br>Investigation(Nom) Renuka-Acc please-Pst<br>Investigation pleased Renuka | രേണുക അന്വേഷണത്തെ ആനന്ദിപ്പിച്ചു}<br>Renuka anweshanathe aanadhippichu<br>Renuka(Nom) investigation-Acc please-Pst<br>Renuka pleased investigation |
| അപകടം കുമാറിനെ സുഖിപ്പിച്ചു<br>Apakadam kumarine sukhippichu<br>Danger(Nom) Kumar-Acc please-Pst<br>Danger pleased Kumar. | കുമാർ അപകടത്തെ സുഖിപ്പിച്ചു<br>Kumar apakadathe sukhippichu.<br>Kumar(Nom) danger-Acc please-Pst<br>Kumar pleased accident |
| അന്വേഷണം ഇന്ദിരയെ സുഖിപ്പിച്ചു<br>Anweshanam indiraye sukhippichu.<br>Investigation(Nom) Indira-Acc please-Pst<br>Investigation pleased Indira | ഇന്ദിര അന്വേഷണത്തെ സുഖിപ്പിച്ചു<br>Indira anweshanthe sukhippichu.<br>Indira(Nom) investigation-Acc please-Pst<br>Indira pleased investigation |
| ഇരുട്ട് ഉസ്മാനെ പ്രസാദിപ്പിച്ചു<br>Irutt usmane prasadhippichu<br>Darkness(Nom) Usman-Acc delight-Pst<br>The darkness delighted Usman. | ഉസ്മാൻ ഇരുട്ടിനെ പ്രസാദിപ്പിച്ചു<br>Usman iruttine prasadhippichu.<br>Usman(Nom) darkeness-Acc delight-Pst<br>Usman delighted darkness. |
| സംഗീതം ഐശ്വര്യയെ പ്രസാദിപ്പിച്ചു<br>Sangeetham Aishwaryaye prasadhippichu<br>Music(Nom) Aishwarya-Acc delight-Pst<br>Music delighted Aishwarya | ഐശ്വര്യ സംഗീതത്തെ പ്രസാദിപ്പിച്ചു<br>Aishwarya sanggethathe prasadhippichu.<br>Aishawarya(Nom) Sangeetham-Acc delight-Pst<br>Aishwarya delighted music |
| ഇരുട്ട് കണ്ണനെ വിഷമിപ്പിച്ചു<br>Irutt Kannane vishamippichu<br>Darkeness(Nom) Kannan-Acc worry-Pst<br>The darkness worried Kannan. | കണ്ണൻ ഇരുട്ടിനെ വിഷമിപ്പിച്ചു<br>Kannan iruttine vishamippichu.<br>Kannan(Nom) darkeness-Acc worry-Pst<br>The darkness made Kannan anxious. |

| | |
|---|---|
| സംഗീതം കാവേരിയെ വിഷമിപ്പിച്ചു<br>Sangeetham Kaveriye vishamippichu.<br>Music(Nom) Kaveri-Acc worry-Pst<br>Music worried kaveri | കാവേരി സംഗീതത്തെ വിഷമിപ്പിച്ചു<br>Kaveri sangeethathe vishamippichu<br>Kaver(Nom) music-Acc worry-Pst<br>Kaveri worried music |
| ജീവിതം ആദർഷിനെ ഉല്ലസിപ്പിച്ചു<br>Jeevitham aadharshine ullasippichu<br>Life(Nom) Aadharsh-Acc please-Pst<br>Life pleased Aadharsh. | ആദർഷ് ജീവിതത്തെ ഉല്ലസിപ്പിച്ചു<br>Aadharsh jeevithathe ullasippichu<br>Aadharsh(Nom) life-Acc delight-Pst<br>Life delighted Aadharsh. |
| മരണം ഇന്ദിരയെ ഉല്ലസിപ്പിച്ചു<br>Maranam Indiraye ullasippichu.<br>Death(Nom) Indira-Acc please-Pst<br>Death pleased Indira | ഇന്ദിര മരണത്തെ ഉല്ലസിപ്പിച്ചു<br>Indira maranathe ullasippichu.<br>Indira(Nom) death-Acc please-Pst<br>Indira pleased Death |
| യാത്ര കലേഷിനെ പേടിപ്പിച്ചു<br>yaathra Kaleshine pedippichu<br>Travel(Nom) Kalesh-Acc make.fear-Pst<br>Travel frightened Kalesh. | കലേഷ് യാത്രയെ പേടിപ്പിച്ചു<br>Kalesh yaathraye pedippichu<br>Kalesh(Nom) Travel-Acc make.fear.Pst<br>Kalesh frightened travel |
| മരണം കല്യാണിയെ പേടിപ്പിച്ചു<br>Maranam Kalyaniye pedippichu.<br>Death frightened Kalyani | കല്യാണി മരണത്തെ പേടിപ്പിച്ചു<br>Kalyani maranthe pedippichu<br>Kalyani frightened death |
| ജീവിതം ആഷിഖിനെ വിനോദിപ്പിച്ചു<br>Jeevitham Aashiqine vinodhippichu.<br>Life(Nom) Aashique-Acc please-Pst<br>Life pleased Aashique | ആഷിഖ് ജീവിതത്തെ വിനോദിപ്പിച്ചു<br>Aashiqine jeevithathe vinodhippichu<br>Aashique(Nom) life-Acc please-Pst<br>Aashique pleased life. |
| യാത്ര ആരതിയെ വിനോദിപ്പിച്ചു<br>Yaathra Aarathiye vinodhippichu<br>Travel(Nom) Aarathy-Acc please-Pst<br>Travel pleased Arathy. | ആരതി യാത്രയെ വിനോദിപ്പിച്ചു<br>Aarathy yathraye vinodhippichu<br>Aarathy(Nom) travel-Acc please-Pst<br>Aarathy pleased travel |
| കളരി അനിലയെ ആഹ്ലാദിപ്പിച്ചു<br>Kalari Anilaye aahladhippichu<br>Kalari(Nom) Anila-Acc please-Pst | അനില കളരിയെ ആഹ്ലാദിപ്പിച്ചു<br>Anila Kalariye ahaldhippichu<br>Anila(Nom) Kalari-Acc please-Pst |

| | |
|---|---|
| Kalari pleased Anila | Anila pleased Kalari |
| കളരി രൂപേഷിനെ രസിപ്പിച്ചു<br>Kalari Roopeshine rasippichu<br>Kalari(Nom) Roopesh-Acc please-Pst<br>Kalari pleased Roopesh | രൂപേഷ് കളരിയെ രസിപ്പിച്ചു<br>Roopesh kalariye rasippichu<br>Roopesh(Nom) Kalari-Acc please-Pst<br>Roopesh pleased kalari |
| നൃത്തം വിജയിയെ മുഷിപ്പിച്ചു<br>Nritham Vijayiye mushippichu<br>Dance(Nom) Vijay-Acc annoy-Pst<br>The dance annoyed Vijay. | വിജയ് നൃത്തത്തെ മുഷിപ്പിച്ചു<br>Vijay nrithathe mushippichu.<br>Vijay(Nom) dance-Acc annoy-Pst<br>Vijay disturbed dance. |
| നൃത്തം നിമിഷയെ ആനന്ദിപ്പിച്ചു<br>Nritham Nimishaye aanadhippichu<br>Dance(Nom) Nimisha-Acc please-Pst<br>The dance pleased Nimisha | നിമിഷ നൃത്തത്തെ ആനന്ദിപ്പിച്ചു<br>Nimisha nrithathe aanadhippichu<br>Nimisha(Nom) dance-Acc please-Pst<br>Nimisha pleased dance |
| ചെണ്ട റുബീനയെ സുഖിപ്പിച്ചു<br>Chenda Rubeenaye sukhippichu<br>Drums(Nom) Rubeena-Acc please-Pst<br>Drums pleased Rubeena | റുബീന ചെണ്ടയെ സുഖിപ്പിച്ചു<br>Rubeena chendaye sukhippichu<br>Rubeena(Nom) drums-Acc please-Pst<br>Rubeena pleased drums |
| ചെണ്ട ജീവനെ പ്രസാദിപ്പിച്ചു<br>Chenda Jeevane prasadhippichu.<br>Drums(Nom) Jeevan-Acc please-Pst<br>Drums pleased Jeevan | ജീവൻ ചെണ്ടയെ പ്രസാദിപ്പിച്ചു<br>Jeevan chendaye prasadhippichu.<br>Jeevan(Nom) drums-Acc please-Pst<br>Jeevan pleased drums |
| ഉത്സവം മനോജിനെ വിഷമിപ്പിച്ചു<br>Ulsavam Manojine vishamippichu<br>Celebration(Nom) Manoj-Acc worry-Pst<br>Celebration worried Manoj. | മനോജ് ഉത്സവത്തെ വിഷമിപ്പിച്ചു<br>Manoj ulsavathe vishamippichu<br>Manoj(Nom) Celebration-Acc worry-Pst<br>Manoj worried Celeberation. |
| ഉത്സവം സുബീനയെ ഉല്ലസിപ്പിച്ചു<br>Utsavam Subeenaye ullasippichu<br>Celebration(Nom) Subeena-Acc please-Pst<br>Celebration pleased Subeena | സുബീന ഉത്സവത്തെ ഉല്ലസിപ്പിച്ചു<br>Subeena utsavathe ullasippichu<br>Subeena(Nom) celebration-Acc please-Pst<br>Subeena pleased Celebration |
| വ്യായാമം ഗ്രീഷ്മയെ പേടിപ്പിച്ചു | ഗ്രീഷ്മ വ്യായാമത്തെ പേടിപ്പിച്ചു |

| | |
|---|---|
| Vyayam Greeshmaye pedippichu<br>Exercise(Nom) Greeshma-Acc made.fear-Pst<br>Exercise frightened Greeshma | Greeshma vyayamathe pedippichu.<br>Greeshma(Nom) exercise-Acc made.fear-Pst<br>Greeshma frightened Exercise |
| വ്യായാമം ജ്യോതിഷിനെ വിനോദിപ്പിച്ചു<br>Vyayam Jyothishine vinodhippichu<br>Exercise(Nom) Jyothish-Acc please-Pst<br>Exercise pleased Jyothish | ജ്യോതിഷ് വ്യായാമത്തെ വിനോദിപ്പിച്ചു<br>Jyothish vyayamathe vindodhippichu<br>Jyothish(Nom) exercise-Acc please-Pst<br>Jyothish pleased Exercise |

### A2: ERPs at the sentence-initial subject noun

Figure. S1 shows the ERPs at the position of the sentence-initial subject noun collapsed over verb type, since the verb is yet to unfold following the nouns. A linear mixed effects model m1 with maximal random effects specification for the single-trial ERP amplitudes at the position of the Np1 was computed in three time windows 300-500ms, 550-750 ms, 750-950 ms. The analysis code and full model outputs are available as R notebooks in the analysis repository online. The analysis revealed no effects.

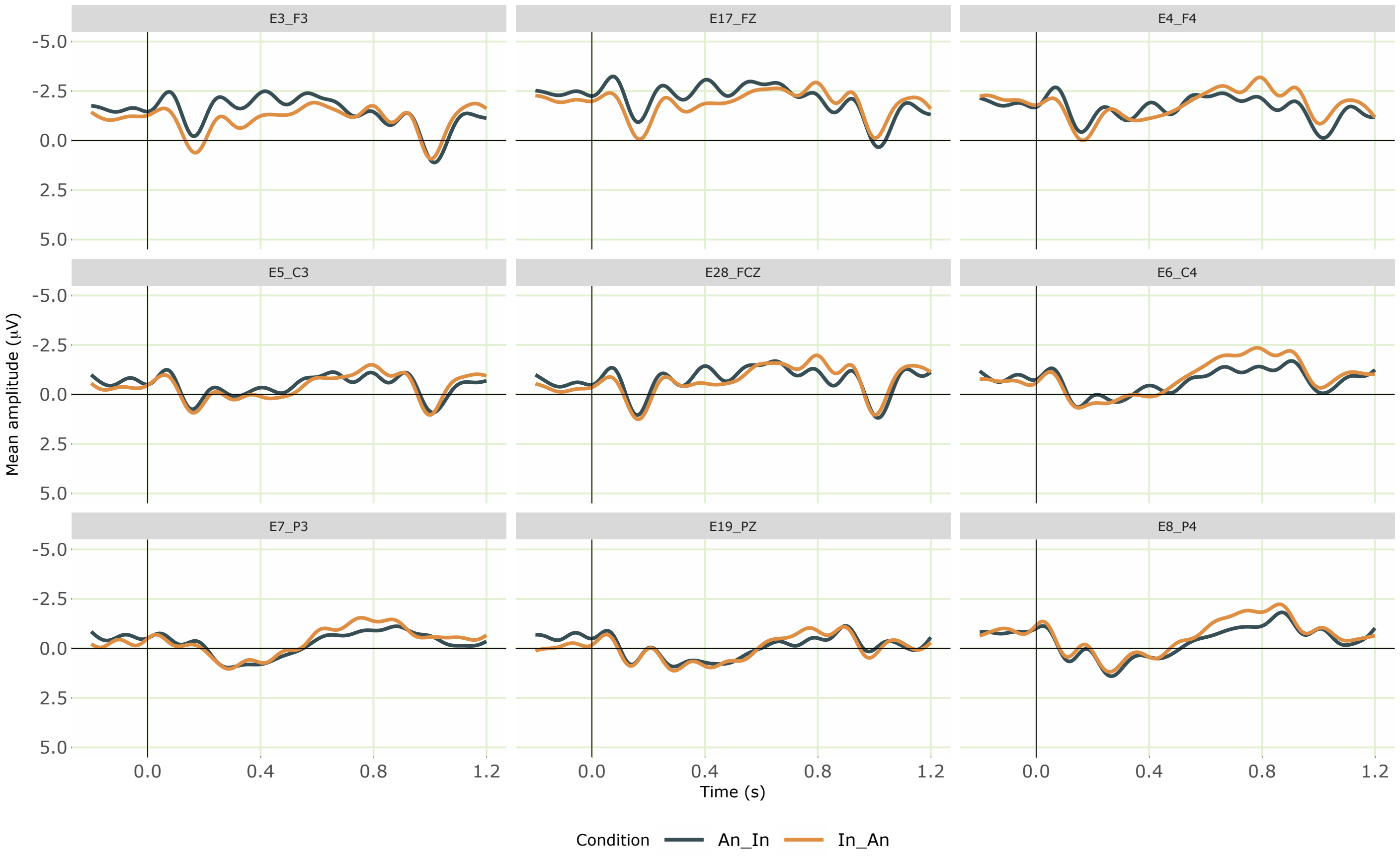


**Figure S1.** Grand averaged ERPs at the nominative animate and nominative inanimate sentence-initial subject nouns from 30 participants (collapsed over verb type, since the verb is yet to unfold following the noun). Negativity is plotted upwards; the time axis runs from -0.2 s to 1.2 s (i.e., -200 ms to 1200 ms) with 0 being the onset of the noun. The dark blue line shows the ERPs for the nominative animate subject and the orange line shows the ERPs for the nominative inanimate subject.

### A3: ERPs at the sentence-medial object noun

Figure. S2 shows the ERPs at the position of the sentence-medial object noun collapsed over verb type, since the verb is yet to unfold following the noun. A linear mixed effects model m1 with maximal random effects specification for the single-trial ERP amplitudes at the position of the Np2 was computed in three time windows 300-500ms, 550-750 ms, 750-950. The analysis code and full model outputs are available as R notebooks in the analysis repository online. The analysis in the 300–500 ms time window yielded informative effects. Type II Wald Chi-squared tests on the fitted model (AIC = 402870.40) in the 300-500 ms time window showed an interaction effect of N1N2Animacy x ROI ($\chi^2(1) = 15.92$, $p = 0.02$, $s = 5.27$). Estimated marginal means on the response scale were computed on the model using the emmeans package (Lenth Russell, 2021) to resolve this interaction. The pairwise contrasts of estimates for N1N2Animacy within each level of ROI revealed a simple effect of N1N2 animacy in the left-frontal region (estimate = -0.808, SE = 0.442, $p = 0.06$, $s = 3.89$). In this region the estimate for animate accusative objects when preceded by inanimate subjects was more negative than (estimate = -1.801, SE = 0.340, $p < 0.001$, $s = 22.95$) inanimate accusative object when preceded by animate subject (estimate = -0.992, SE = 369, $p = 0.007$, $s = 7.09$).

We interpret that the negativity effect (300-500 ms) observed in the left-frontal region for animate objects preceded by inanimate subjects as reflecting mismatch between different prominence hierarchies (Bornkessel-Schlesewsky and Schlesewsky, 2009). In this configuration, the object position and accusative case mark the animate argument as an undergoer, while its animacy renders it a better candidate for the actor role. When the parser encounters this animate argument in object position, the conflict between positional and case cues and animacy-based expectations cleads to an N400 effect.

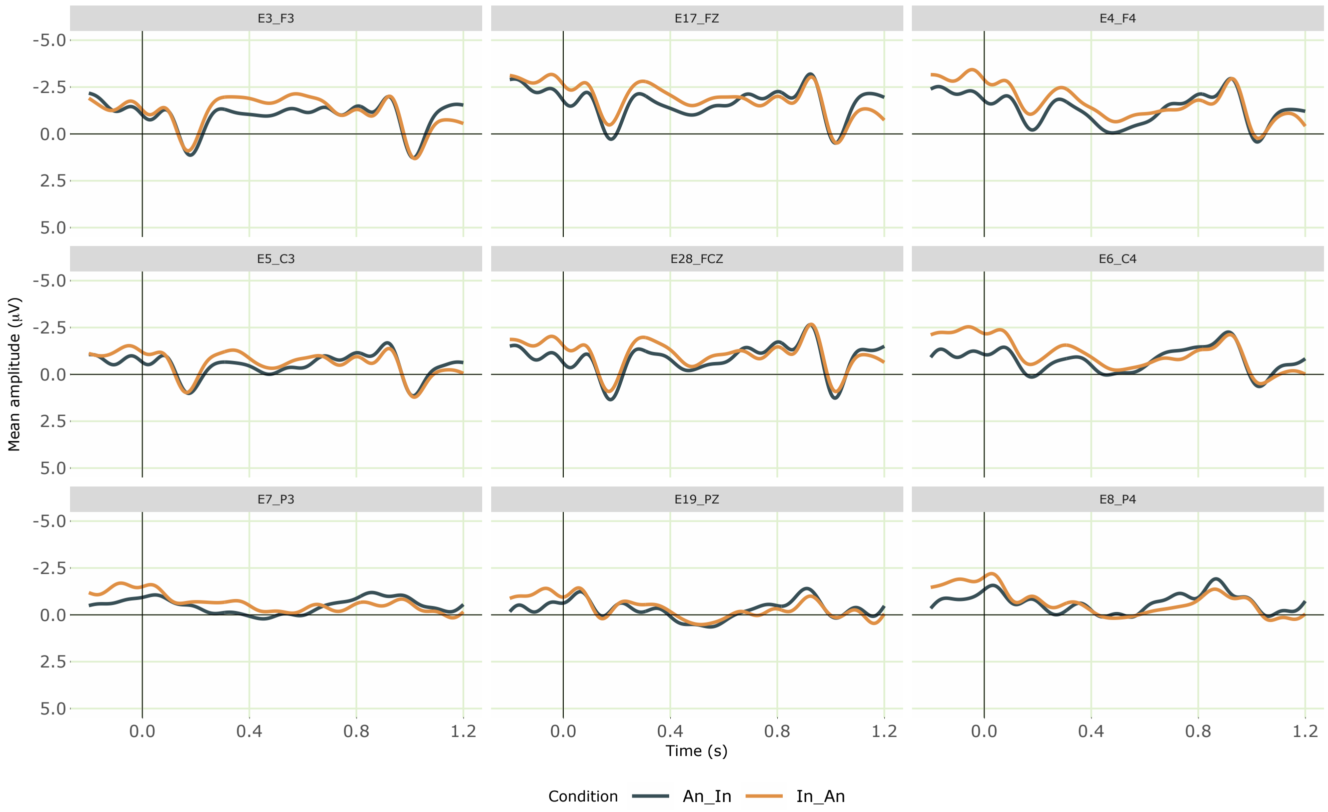


**Figure S2.** Grand averaged ERPs at the accusative inanimate and accusative animate sentence-medial object nouns from 30 participants (collapsed over verb type, since the verb is yet to unfold following the noun). Negativity is plotted upwards; the time axis runs from -0.2 s to 1.2 s (i.e., -200 ms to 1200 ms) with 0 being the onset of the noun. The dark blue line shows the ERPs for the accusative inanimate object preceded by animate subject and the orange line shows the ERPs for the accusative animate object preceded by inanimate subject.

### A4: ERPs at the Sentence-final Verb

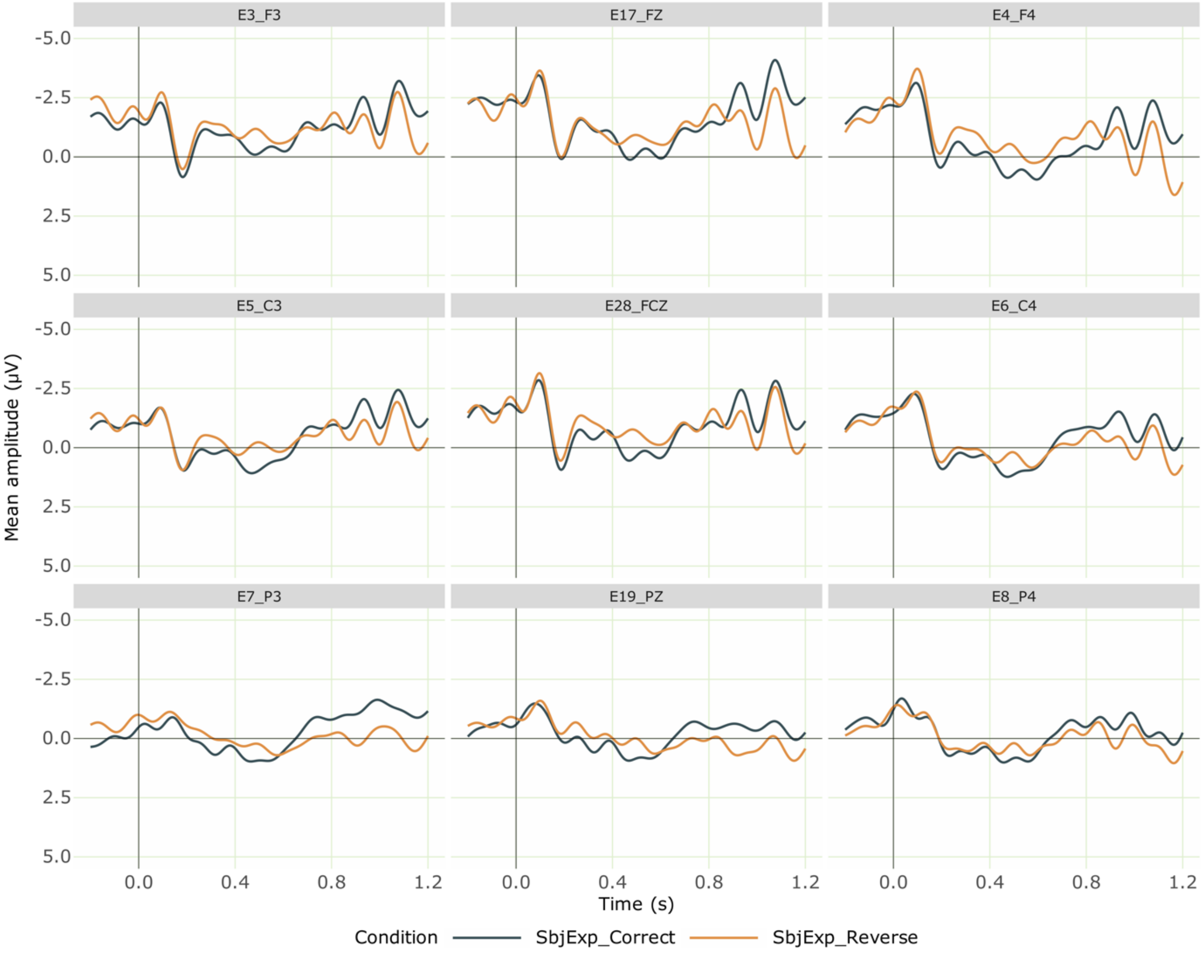


**Figure S3.** Grand averaged ERPs at the verb for the critical conditions from 30 participants. Negativity is plotted upwards; the time axis runs from -0.2 s to 1.2 s (i.e., -200 ms to 1200 ms) with 0 being the onset of the critical verb. The dark blue line represents the correct SubjExp condition and the orange line represents the role reversed SubjExp condition, which elicited a centro-parietal negativity followed by a positivity effect.

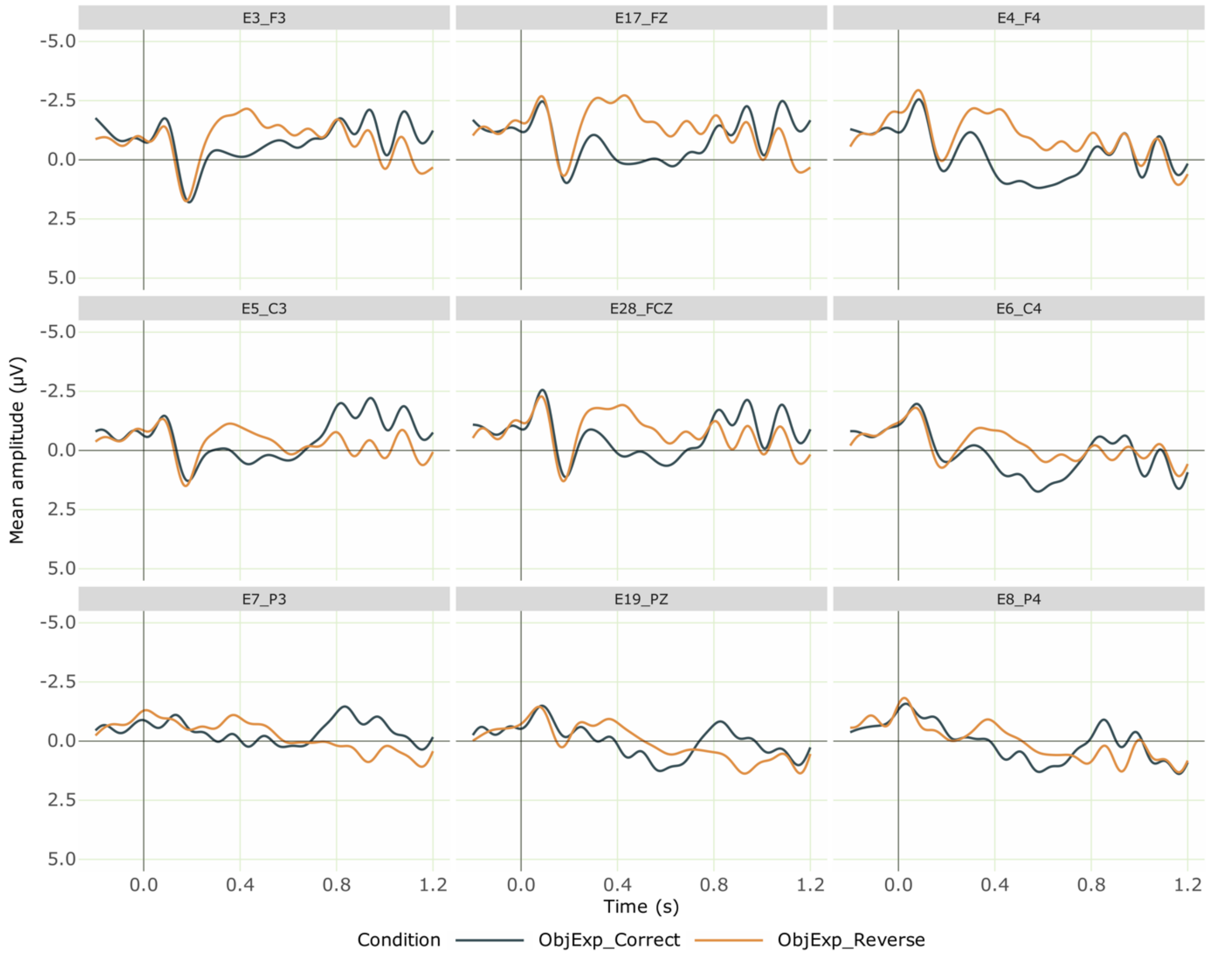


**Figure S4:** Grand averaged ERPs at the verb for the critical conditions from 30 participants. Negativity is plotted upwards; the time axis runs from -0.2 s to 1.2 s (i.e., -200 ms to 1200 ms) with 0 being the onset of the critical verb. The dark blue line represents the correct ObjExp condition, while the orange line shows the role reversed ObjExp condition, which elicited a centro-parietal negativity followed by a positivity effect.
